# A Hessian-based decomposition characterizes how performance in complex motor skills depends on individual strategy and variability

**DOI:** 10.1101/645317

**Authors:** Paolo Tommasino, Antonella Maselli, Domenico Campolo, Francesco Lacquaniti, Andrea d’Avella

**Affiliations:** Laboratory of Neuromotor Physiology, IRCCS Fondazione Santa Lucia, Rome, Italy; Synergy Lab, Robotics Research Centre, School of Mechanical and Aerospace Engineering, Nanyang Technological University, Singapore, Singapore; Department of Systems Medicine and Center of Space Biomedicine, University of Rome Tor Vergata, Rome, Italy; Department of Biomedical and Dental Sciences and Morphofunctional Imaging, University of Messina, Messina, Italy

## Abstract

In complex real-life motor skills such as unconstrained throwing, performance depends on how accurate is on average the outcome of noisy, high-dimensional, and redundant actions. What characteristics of the action distribution relate to performance and how different individuals select specific action distributions are key questions in motor control. Previous computational approaches have highlighted that variability along the directions of first order derivatives of the action-to-outcome mapping affects performance the most, that different mean actions may be associated to regions of the actions space with different sensitivity to noise, and that action covariation in addition to noise magnitude matters. However, a method to relate individual high-dimensional action distribution and performance is still missing. Here we introduce a decomposition of performance into a small set of indicators that compactly and directly characterize the key performance-related features of the distribution of high-dimensional redundant actions. Central to the method is the observation that, if performance is quantified as a mean score, the Hessian (second order derivatives) of the action-to-score function determines how the noise of the action distribution affects the average score. We can then approximate the mean score as the sum of the score of the mean action and a tolerance-variability index which depends on both Hessian and action covariance. Such index can be expressed as the product of three terms capturing noise magnitude, noise sensitivity, and alignment of the most variable and most noise sensitive directions. We apply this method to the analysis of unconstrained throwing actions by non-expert participants and show that, consistently across four different throwing targets, each participant shows a specific selection of mean action score and tolerance-variability index as well as specific selection of noise magnitude and alignment indicators. Thus, participants with different strategies may display the same performance because they can trade off suboptimal mean action for better tolerance-variability and higher action variability for better alignment with more tolerant directions in action space.

**Author summary:** Why do people differ in their performance of complex motor skills? In many real-life motor tasks achieving a goal requires selecting an appropriate high-dimensional action out of infinitely many goal-equivalent actions. Because of sensorimotor noise, we are unable to execute the exact same movement twice and our performance depends on how accurate we are on average. Thus, to understand why people perform differently we need to characterize how their action distribution relates to their mean task score. While better performance is often associated to smaller variability around a more accurate mean action, performance also depends on the relationship between the directions of highest variability in action space and the directions in which action variability affects the most the outcome of the action. However, characterizing such geometric relationship when actions are high dimensional is challenging. In this work we introduce a method that allows to characterize the key performance-related features of the distribution of high-dimensional actions by a small set of indicators. We can then compare such indicators in different people performing a complex task (such as unconstrained throwing) and directly characterize the most skilled ones but also identify different strategies that distinguish people with similar performance.

## 1 Introduction

In many goal-directed human behaviors, such as throwing a projectile towards the center of a target, performance depends on how accurate the outcome of repeated actions is. In throwing tasks, *accuracy* of a single throw may be quantified by a *score*, e.g. a scalar function that penalizes/rewards motor outcomes depending on their distance from the desired target position [18, 14]. In this perspective, the goal of a thrower would be that of minimizing/maximizing the mean score over repeated trials [26, 8]. Because of bias and noise in the sensorimotor transformations mapping goals into actions [13, 25], the score typically varies across trials and hence the *performance* of a throwing strategy, defined as its mean score, will in general depend on the *distribution* of motor actions [18, 23].

In many tasks, the relationship between motor actions and their outcomes is *redundant* [2] and to different actions there might correspond the same task outcome, hence the same score. As an example, consider the throwing task shown in Fig 1A where the outcome space is the two-dimensional space of all the possible landing positions of the ball on a vertical board, and the score depends on where the ball lands with respect to the aimed target. The landing position of the ball ultimately only depends on the position and velocity with which the ball is released (*actions*). The center of the target then can be hit with different combinations (or *covariations*) of such action variables: for instance one can hit the target by releasing the ball from different positions modulating the velocity vector accordingly, or from a fixed position but with different combinations of vertical and horizontal velocities. These different but *task-equivalent* actions resulting in the same landing position form a subset of the action space which is called *solution manifold*. Key questions in human motor control are then whether different individuals select specific action distributions to achieve a given performance level, what characteristics of the action distribution relate to performance, and how action distributions change when performance improves with practice [2, 9, 24, 18, 30, 10].

**Figure 1.**
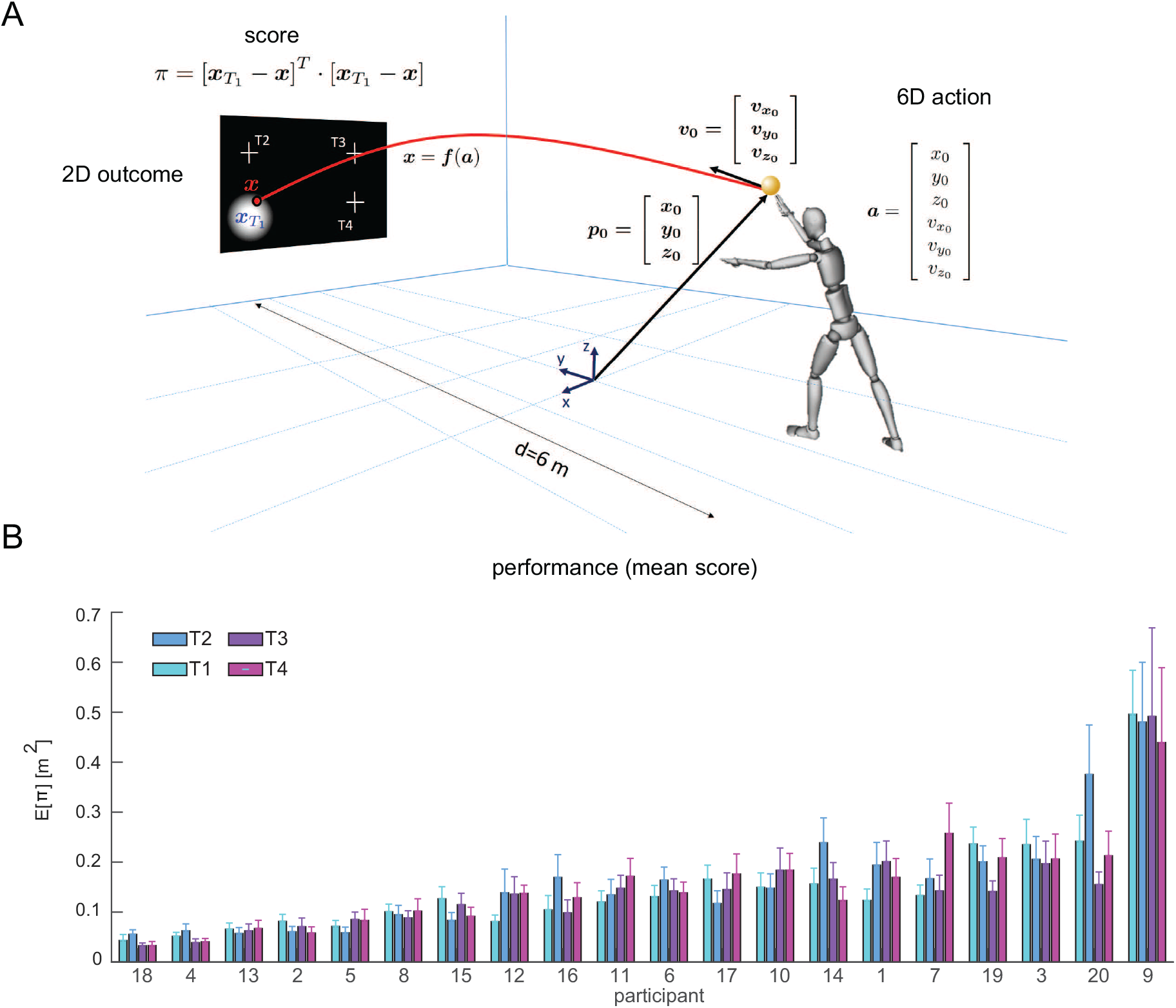
Unconstrained throwing task and experimental performance. (A) Schematic representation of the unconstrained overarm throwing task considered in this study as an example of a complex motor skill. The release parameters characterizing a throwing action are six-dimensional (three position components and three velocity components). (B) Performance (mean squared error ± SE) across participants (n = 20) and four targets.

In well-developed motor skills, outcomes are typically unbiased (zero mean error) and hence outcome variability (or precision) is usually taken as a measure of performance. Indeed, when learning a new motor skill involving redundant actions with different amounts of noise or noise tolerance in different regions of the action space, participants improve performance by selecting actions whose outcome is less affected by motor noise [31, 33]. In this perspective, the relationship between action distribution and outcome variability in goal-directed behaviors has been addressed by computational approaches that take into account the geometry of the mapping between actions and outcomes, also known as the goal function, near the solution manifold. Methods such as the Uncontrolled Manifold (UCM) [27, 19] and the Goal-Equivalent Manifold (GEM)[6, 7] typically approximate the (non-linear) action-to-outcome function with a (locally) linear map, which relates (stochastic) perturbations of the mean action to the precision (variance or covariance) of task outcomes. More specifically, the gradient (or Jacobian) of such mapping is employed to quantify action variability along *task-irrelevant* directions (directions parallel to the solution manifold) and *task-relevant* directions (directions orthogonal to the solution manifold). The UCM applied to reaching, pointing and throwing tasks [27, 19, 35] has shown that *covariation* between redundant actions is an important mechanism used by skilled performers to push motor variability along the task-irrelevant directions, hence increasing the precision of their task outcomes. Differently from UCM, which only quantifies motor strategies in terms of the *alignment* between action variability and task-relevant/task-irrelevant directions, the GEM approach takes also into account the *sensitivity* of the solution manifold to local perturbations. Then, different choices of mean actions may result in different amounts of outcome variability because of the specific alignment and sensitivity to the different mean actions, i.e. factors depending on the local geometry of the goal function, rather than only on different amounts of action variability.

The impact of the interplay between motor variability and task geometry on performance in goal-directed behaviors has been also investigated with an approach that needs no assumption on the smoothness of the action-to-score mapping, but relies on the computation of surrogate data or on the exhaustive search of optimal data distributions, rather than analytic descriptions, to explore its geometry [23, 5, 30]. The Tolerance-Noise-Covariation (TNC) method [23] allows to quantify the difference in performance between two series of trials as the sum of three contributions. The *tolerance* component is associated with potential differences in the *sensitivity* of the local action-to-score mapping geometry associated with different choices of the mean action. The *noise* component quantifies the impact of different amount of action variability in the two series. Finally, the *covariation* component accounts for the impact of different alignment of the action variability with the local geometry. A revised version of the method (TNC-Cost analysis [5]) allows to assess the three components of a single series of trials with respect to optimal performance.

A key aspect of the TNC approach that makes it particularly suitable for the analysis of inter-individual differences in goal-directed behaviors is its focus on the relationship between action distribution and performance as mean score. While the UCM and GEM approaches focus on variability in action space and outcome space, the TNC decomposition shows how the mean score depends on the choice of the mean action in addition to variability. Furthermore, by identifying different contributions to the mean score, the TNC approach allows to characterize individual performances with a higher level of details: for instance, a participant could perform the task with a higher variability than a peer, but achieve the same level of performance thanks to a better alignment. However, because the computations of the costs depend on numerical procedures that become cumbersome for high dimensional action spaces [30], the TNC analysis has been applied only to simple tasks, such as a planar virtual version of the skittles game in which one can control the angle and velocity of ball release [23, 36].

In the present work, our goal was to characterize inter-individual differences in the relationship between action-variability and performance in a complex, real-life motor skill. We asked twenty non-expert participants to perform unconstrained overarm throws at four different targets placed on a vertical plane at 6 m distance, as depicted in Fig 1A. When we analyzed the time-course of the whole-body kinematics during the throwing motion [21, 20], we found that throwing styles, i.e. the average trajectory of each limb segment, differed considerably across individuals. Similarly, we found large differences in individual performances, as shown in Fig 1B. Here, we focused on inter-individual differences in terms of release parameters distribution and throwing performance, as differences in throwing styles may translate into different release strategies, which may or may not correspond to differences in the mean score. We aimed at characterizing the key performance-related features of the individual release parameters distribution. We also aimed at identifying consistent features that could explain why participants differed in performance and how they could achieve similar performance levels with different strategies.

To identify the different contributions of the distribution of throwing actions to performance, we introduced a novel analytic method that can be applied to an unconstrained throwing task, described by at least six release parameters, overcoming the computational limitation of the TNC method, which requires a number of numerical operations that scale exponentially with the number of dimensions of the action space. Our approach is based on second order derivatives of the action-to-score function and depends on the following two assumptions: i) the action distribution is sufficiently localized in a region of the action space; ii) the score function, although non-linear, can be adequately approximated with a second-order Taylor-expansion. Hence, we make use of the Hessian matrix, i.e., the matrix of second-order partial derivatives, rather than the Jacobian, to estimate the local tolerance of the score as well as to estimate the alignment of action covariance with the curvature of the action-to-score mapping.

## 2 Results

To relate performance in a goal-directed motor task to the distributions of motor actions, we introduce a decomposition of the mean score in terms of a few parameters depending on the mean action, the Hessian of the action-to-score mapping computed at the mean action, and the covariance of the action distribution. Fig 2A illustrates the variables and functions describing the relationship between an action (*a*) and a score (or loss, *π*) assigned to the outcome (*x*) of the action. Such score represents a scalar measure of inaccuracy of the outcome with respect to the goal (or target, *x*_*T*_) and, thus, it is a composite function of the action-to-outcome mapping (*x* = *f* (*a*)) and the outcome-to-score mapping (*π* = *s*^*a*^(***x***; ***x***_*T*_)).

**Figure 2.**
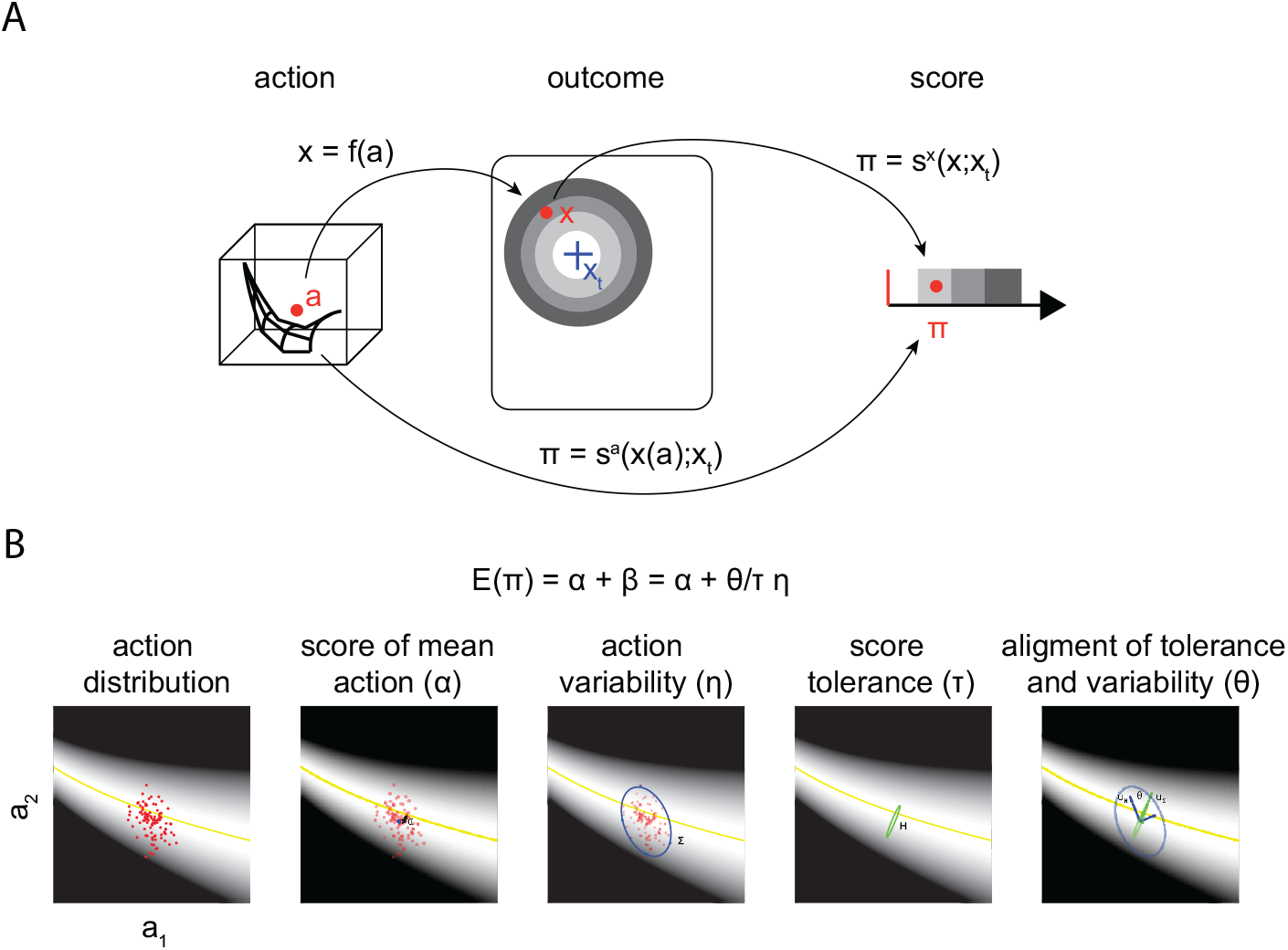
Definition of variables (action, outcome, and score) and schematic overview of the proposed approach to decompose the mean score (performance). (A) An action, represented by a vector *a* in a high-dimensional *action space* (illustrated by a *red* circular marker in a three-dimensional action space), leads to an outcome, ***x*** = *f* (***a***), i.e. a vector typically represented in *outcome* or *goal space* with less dimensions. The outcome is then associated with a corresponding score, *π* = *s*^*x*^(***x, x***_*T*_), based on its relation with respect to the aimed target. The set of points in action space associated with an outcome achieving the goal constitute the *solution manifold* (illustrated by the black mesh). (B) Illustration of the Hessian-based decomposition of the mean score in the case of two-dimensional actions. The action-to-score mapping is represented by the gray-shaded areas. A set of actions (small red markers, *first panel*) has a distribution characterized by the mean (large blue marker, *second panel*) and covariance (Σ, blue ellipse in the *third and fifth panels*). The tolerance of the score to variations of the actions around their means is characterized by the Hessian (*H*, green ellipse in the *fourth panel*). The mean score (*E*(*π*)) is decomposed as the sum of the score of the mean action (*α*) and a *tolerance-variability* index (*β*) expressed as the product of three terms: the reciprocal of the *tolerance* (*τ*), the *uncorrelated noise* (*η*), and the *alignment* (*θ*), i.e. a scalar characterizing whether the most sensitive directions of the Hessian (green arrows ***u***_*H*_, *fifth panel*) are aligned to the directions of highest action variability (blue arrows ***u***_Σ_).

If we consider an action distribution with mean ***ā*** and covariance Σ^*a*^, by using a second-order Taylor-expansion of the action-to-score function around the mean action (see sec:Methods), the mean score can be approximated as:

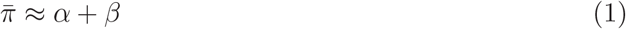

where

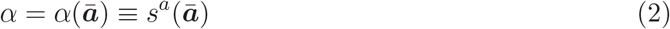

is *the score of the average action*, and

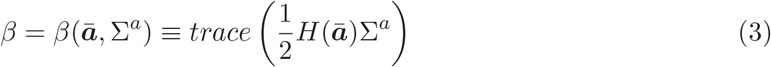

is a *tolerance-variability index* which captures how local non-linearities of the action score (the Hessian *H*(***ā***)) and variability in the action strategy (Σ^*a*^) affect performance.

Furthermore, by defining *score sensitivity* as the trace of *H* and *score tolerance* as its reciprocal

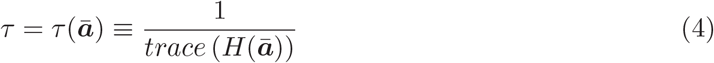

and *uncorrelated noise* as the trace of the covariance matrix (i.e., the total variation of the action distribution, providing a scalar measure of the overall amount of action variability)

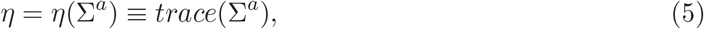

we can express the *tolerance-variability index* as

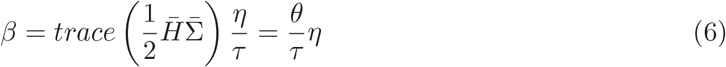

where we introduced the *alignment* as

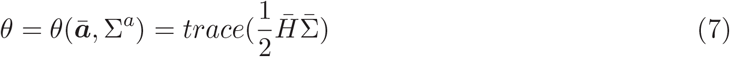

with 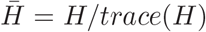 and 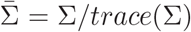. The alignment is here a scalar that measures the effect on performance of the relative orientation between the directions of highest sensitivity of the action-to-score mapping with respect to the directions of highest variability of the actions (see Methods).

Fig 2B provides a graphical illustration of the decomposition approach in the case of two-dimensional actions, as in a throwing task in which only the horizontal and vertical components of the release velocity can vary. The score associated to each action is indicated by the *gray-shaded* background. The yellow curve at the center of the white area represents the *solution manifold*, the set of actions that accurately achieve the goal. The distribution of actions is characterized by their mean (***ā***) and covariance (Σ) while the local geometry of the action-to-score mapping is characterized by the Hessian (*H*) computed at the mean action.

In sum, the expected value of the score (*E*(*π*)) can be decomposed as the sum of the score of the mean action (*α*) and the tolerance-variability index (*β*)

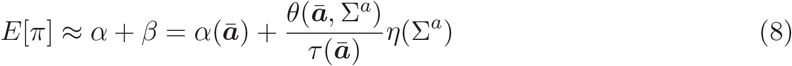

where *β* is determined by the interplay between the sensitivity of the performance to action variability (reciprocal of the tolerance *τ*), the magnitude of the action variability *η*, the alignment between the Hessian and the action covariance *θ*.

### 2.1 Simulated 2D throwing examples

To illustrate how our decomposition allows to identify the key features in the action distribution contributing to performance, we consider a toy model of a throwing task in which a projectile is released from a fixed position with given horizontal and vertical velocity components and must hit a target at a fixed height on a vertical plane at a fixed distance. Additionally, we assume that the horizontal component of the release velocity lies in the vertical plane containing the release position and the target, so that the action *a* is a two-dimensional (2D) vector, the projectile trajectory lies on a vertical plane, and the outcome (*x*, i.e., the arrival height) is a scalar (Fig 3A). The score *π*(*x*; *x*_*T*_) is the squared distance of the arrival position from the target.

**Figure 3.**
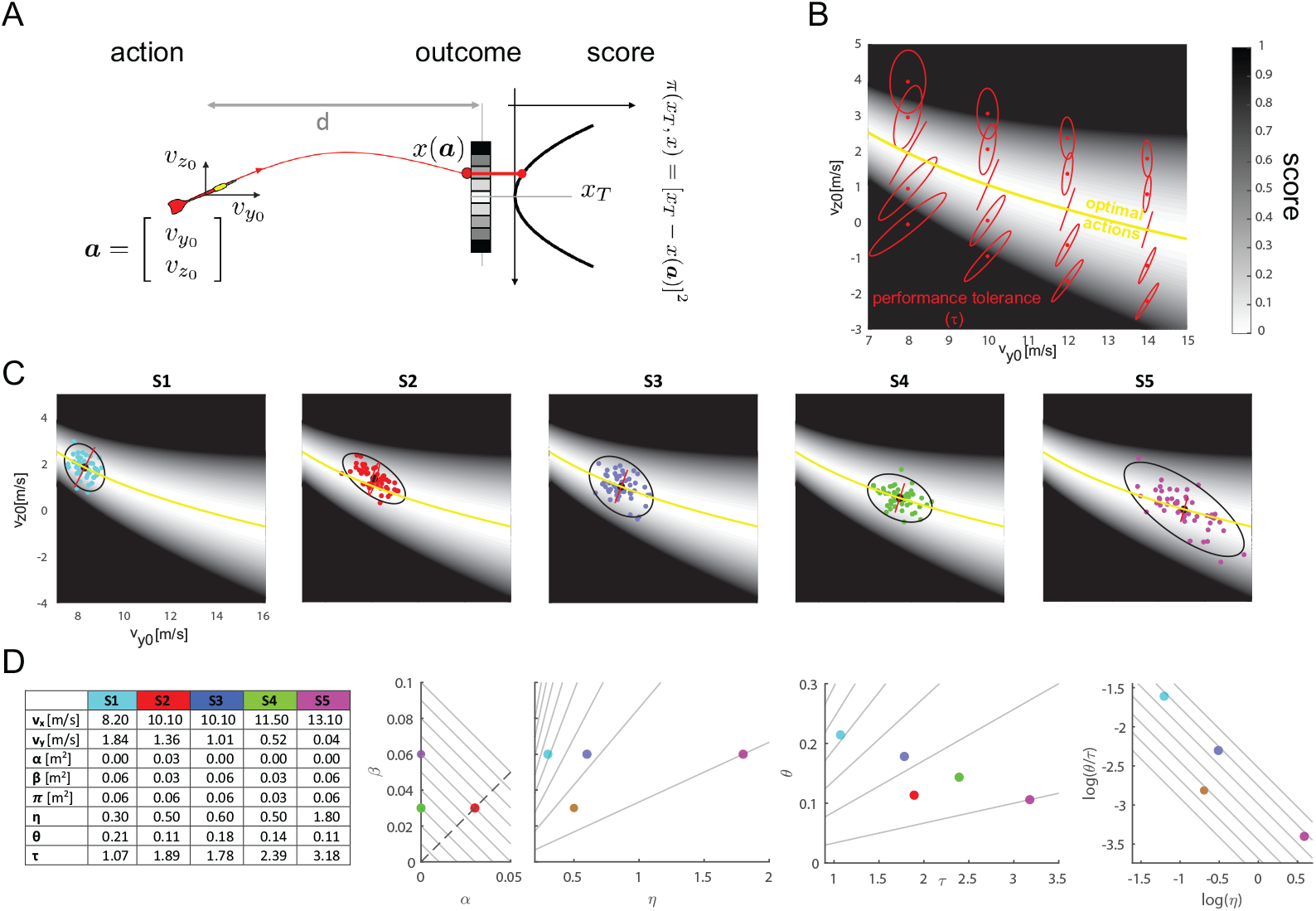
Examples of different throwing strategies in a simulated 2D throwing task. (A) Toy model of a throwing task in which the action (***a***) is characterized by only two parameters, i.e. the horizontal (*v*_*y*0_) and the vertical (*v*_*z*0_) release velocity components, the outcome (*x*) is the arrival position of the projectile on a vertical plane at a distance *d* from the release position, and the score (*π*) is the squared distance of the arrival position from a target (*x*_*T*_). (B) Action score (gray-shaded background), solution manifold (yellow line), and Hessian (red ellipses) in a region of release parameters. The score sensitivity for different throwing strategies, is captured by the Hessian, which is represented with red ellipses whose major axis represents, locally, the direction of maximum sensitivity (smaller noise tolerance). (C) Five (simulated) individual strategies. For each strategy (*S*1-*S*5, *different panels and colors*) the individual throwing actions are shown together with mean (*large black circle*), their covariance (*black ellipse*, 95% C.L.) and the Hessian at the mean action (*red ellipse*). (D) Hessian-based decomposition of the mean score of the five strategies. For each strategy, the mean release parameters (*v*_*y*0_ and *v*_*z*0_)), the mean score 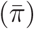, and the indicators from the decomposition (score of the mean action *α, tolerance-variability* index *β, tolerance* (*τ*), *uncorrelated noise η*, and *alignment θ*) are presented in the table. The four scatter plots illustrate how these indicators allow to differentiate strategies with different performance as well as strategies with same performance. See text for more details.

As in Fig 2B, since the actions are 2D, in Fig 3B we can represent the action-to-score function as contours in the action plane (i.e., the plane *v*_*x*0_-*v*_*z*0_ of the horizontal and vertical release velocities) with gray shading (lighter shades for lower error) and the solution manifold, i.e. the set of optimal actions which results in 0 penalty score, with a yellow curve. Fig 3B also shows the local sensitivity of the score, i.e. the key element of our decomposition, which is captured by the Hessian of the action-to-score function. For different values of the mean action (i.e. 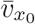 and 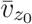), the sensitivity of the mean score is represented by a red ellipse. The longer axis of the ellipse, whose length is proportional to the largest eigenvalues of the Hessian matrix (see Methods), corresponds to the direction in which a given deviation of an action from the mean affects the mean score the most, i.e. the direction of maximal sensitivity or minimal tolerance to noise. The length of each axes of the ellipse indicates how much a deviation in that direction affects the mean score. Consequently, action distributions with the same amount of variability (uncorrelated noise) will have different performances depending on the alignment of their covariance with the direction of maximal sensitivity and on the lengths of the axes of the local sensitivity ellipses. Note that the orientation of the ellipses, their size, and the ratio of their axes depend on the position on the action plane. On the solution manifold, the ellipses have only one axis with non-zero length, indicating that there is only one direction in action space which is relevant for the score. This direction however, is not constant, but changes along the solution manifold. Also note that the ellipses are smaller for larger horizontal velocities 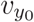, indicating that the score is more tolerant for faster throws.

#### Iso-performing strategies: trade-offs between indicators derived from the Hessian-based decomposition

Fig 3C shows simulated action distributions for five different strategies, each represented by 50 throws with 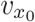 and 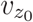 drawn from a bi-dimensional Gaussian distribution with a different mean and a different covariance. The plots are arranged from *left* to *right* according to the magnitude of the mean horizontal velocity 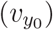. The fourth strategy (*P*4 *green*) is the best performing of the five, as it has a mean action on the solution manifold and the direction of maximal action variability (major axis of the *black* covariance ellipse) is almost orthogonal to the direction of maximal noise sensitivity (major axis of the *red* local sensitivity ellipse). The second strategy (*S*2 *red*) has a worse performance than the fourth, even if the two have the same magnitude of uncorrelated noise *η* and a similar low alignment of the covariance with the local noise sensitivity. This is because its mean action is not on the solution manifold, as it has a vertical release velocity larger than optimal. The remaining three strategies (*S*1 *cyan, S*3 *blue, S*5 *magenta*) have different amount of noise and different levels of alignment with the local noise sensitivity but they have the same performance, also equal to that of *S*2. However, it is difficult to recognize by visual inspection of the action distibution that the four strategies (*S*1, *S*2, *S*3, and *S*5) have the same performance and to characterize their differences.

The Hessian-based decomposition of the mean score of the five strategies according to equations (1) and (6) is illustrated in Fig 3D. Four scatter plots between pairs of the different indicators derived from the decomposition (score of the mean action *α, tolerance-variability* index *β, tolerance τ, uncorrelated noise η*, and *alignment θ*) allow to compactly describe the five strategies, to directly relate them to performance, and to characterize their differences. The *α*-*β* scatter plot shows that strategies with the same performance may trade-off accuracy of the mean action (*α*) with the combined effect of action variability and local geometry of the action-to-score mapping (*β*). The diagonal lines with slope −1 represent *iso-performing strategies* (constant *α* + *β* equal to different values of 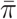). Strategy *S*4 is the best performing one as it has zero *α* and a low value of *β*. Strategy *S*2 has the same low *β* as *S*4 but a higher 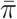 due to non-zero *α. S*2 performance matches that of the remaining three strategies (*S*1, *S*3, and *S*5) as they have higher *β* but null *α* but the four strategies lie on the same diagonal line.

The *η*-*β* scatter plot reveals that *S*1, *S*3, and *S*5 achieve the same *β* with different amounts of noise (*η*). This occurs because their action distributions differ in the alignment of the direction of largest variability with the most noise sensitive directions (*θ*) and because their mean action is located in regions of the action plane with different noise tolerance (*τ*), as shown in the *τ* -*θ* scatter plot. The lines in the *η*-*β* scatter plot represent strategies that have the same slope in the linear dependence of the tolerance-variability index *β* on uncorrelated noise *η*, because they have a given ratio between alignment *θ* and tolerance *τ*, as 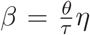. Analogously, the lines in the *τ* -*θ* scatter plot represent strategies with equal 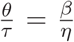. As the horizontal speed increases, the distributions are located in regions with higher tolerance *τ* and they have a lower alignment *θ*.

The trade-off between the effect of variability and the effect of local score sensitivity can be best illustrated in the 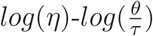 scatter plot. Since 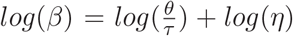, the diagonal lines with slope −1 represent strategies in which the same *β* results from an increase in noise *η* compensated by a decrease in alignment *θ* or an increase in tolerance *τ*. *S*1 (*cyan*) is in a region with lower tolerance (*τ* = 1.07, see table in Fig 3D) and has a higher alignment (*θ* = 0.21), but it compensates for such unfavorable conditions with lower noise (*η* = 0.22). *S*3 (*blue*) has a larger mean horizontal release and a corresponding higher tolerance (*τ* = 1.78) and it achieves the same performance with larger noise (*η* = 0.37). Finally, *S*5 (*magenta*) tolerates even larger noise (*η* = 0.61) by virtue of the higher tolerance (*τ* = 3.18) associated to the largest mean release horizontal velocity.

In sum, the four iso-performing strategies (*S*1, *S*2, *S*3, and *S*5) illustrate two different trade-offs between indicators derived from the Hessian based decomposition. *S*2 vs. *S*1, *S*3, and *S*4 trade-off bias with tolerance-variability. *S*1 vs. *S*3 vs. *S*5 trade-off noise alignment and sensitivity with noise magnitude. In both cases, the indicators allow to compactly characterize the features in the action distribution determining a given performance and those distinguishing different strategies, even when they have the same performance.

#### Comparison with TNC, UCM, and GEM analyses

The 2D throwing toy model also allows to compare the Hessian-based decomposition with existing computational methods that have been introduced for analyzing the relationship between an action distribution and its mean score (TNC), for decomposing action variability into task-relevant and task-irrelevant components (UCM), and for relating action variability and outcome variability (GEM).

The TNC-Cost analysis [5] characterizes an action distribution with three indicators (or costs) computed by comparing the mean score of the distribution with the score of a transformed distribution, optimal according to different criteria. The *tolerance* cost (T-cost) represents the difference between the mean score of the distribution and the mean score of the same distribution translated to an optimal location, i.e. a location with the lowest possible score. For the *noise* cost (N-cost) the optimal distribution is obtained by shrinking the distribution (i.e., reducing the uncorrelated noise *η*) without affecting the mean. The *covariation* cost (C-cost) is computed using an optimal distribution with the same mean and the same individual action components (e.g. 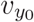 and 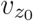 in our 2D throwing example) but with an optimal re-association of those components in different actions, i.e. with an optimal covariation. These indicators are related to the indicators of our decomposition (see Appendix A.2) but they represent a re-combinations of those indicators and, more importantly, do not provide a direct decomposition of the mean score, thus they do not allow to describe the performance trade-offs between indicators presented above. Also, critically, as the computation of both the N-Cost and the C-cost relies on exhaustive searches, they require a number of operations that scales exponentially with the dimensions of the action space and they are not feasible for 6D actions, as in unconstrained throwing.

Fig 4A presents an example of three simulated strategies with the same performance and an *α*-*β* trade-off. Increasing the mean horizontal release velocity 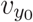, a larger bias *α* is compensated by a lower tolerance-variability index *β* achieved by a lower alignment *θ* and higher tolerance *τ* with constant noise *η*. The T-Cost is constant, as the three distributions have the same mean score and the same covariance matrix, i.e. they differ only for the position of the mean action, and therefore they have the same difference with the mean score of the optimally translated distribution. Thus, the T-Cost does not distinguish iso-performing strategies with different bias when this is compensated by better alignment and tolerance. Moreover, the strategies have different N-cost, even if their action distribution have the same covariance matrix. This is due to the fact that when the action-to-score function is smooth and the dispersion of the actions not too high, as in this case, the optimal distribution used to compute the N-cost has zero noise, i.e. is concentrated on the mean action, and the N-cost corresponds to 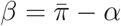.

**Figure 4.**
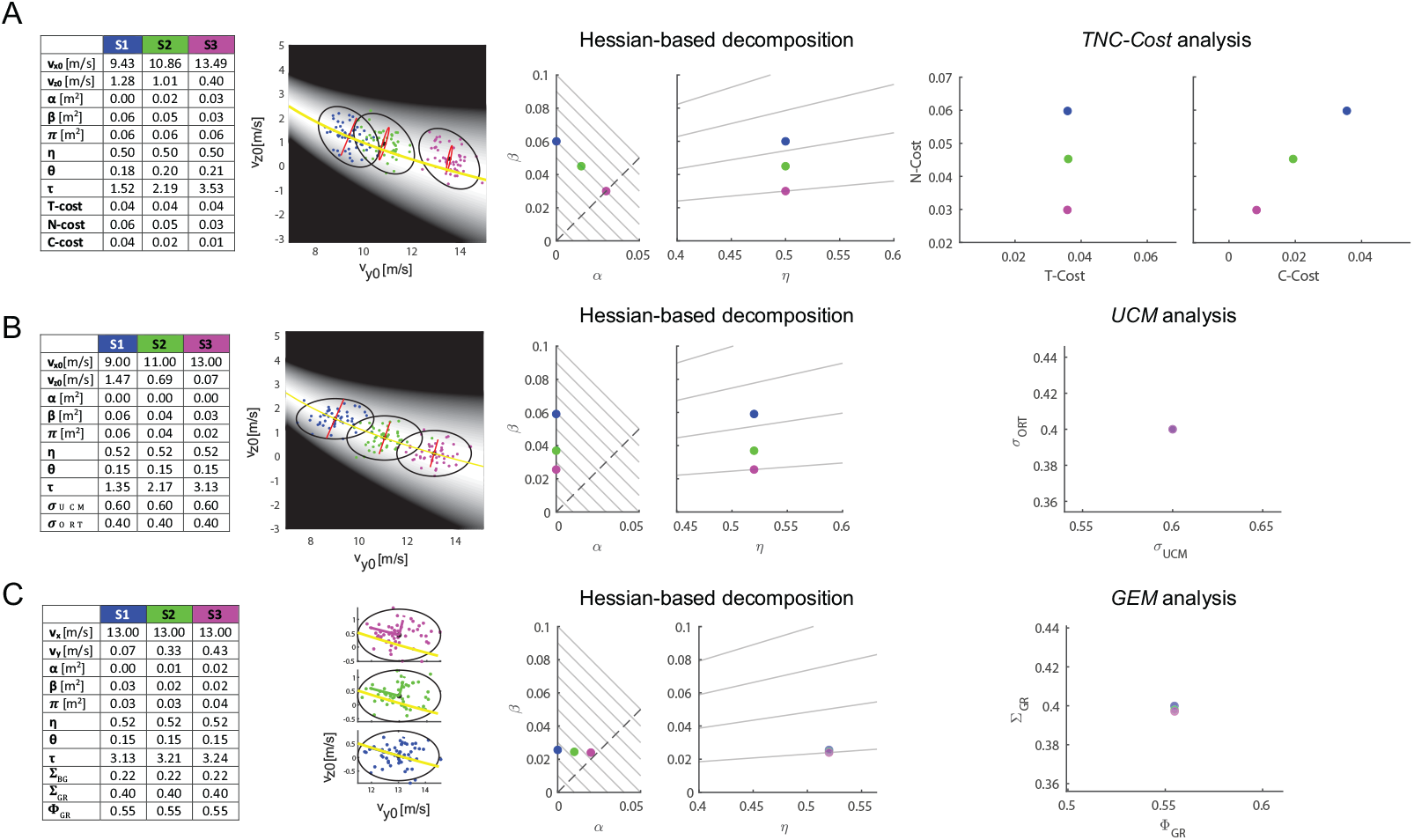
Comparison of the Hessian-based decomposition with existing computational methods for the analysis of throwing strategies in a 2D throwing task. (A) Comparison with TNC-Cost analysis. Three strategies (*S*1 *blue, S*2 *green, S*3 *magenta*) are simulated by generating a distribution of 50 release parameters (*v*_*y*0_,*v*_*z*0_) by randomly sampling from a bi-dimensional Gaussian distribution with a given mean and covariance (see parameters in the table on the *left*). The strategies have the same performance because they trade-off bias (*α*) with tolerance-variability (*β*), as can be clearly observed in the *α*-*β* scatter plot. Moreover, the strategies have equal noise (*η*). However, the strategies have equal T-cost but different N-Cost, which then represent a recombination of the indicators extracted from the Hessian-based decomposition. Moreover, as the TNC costs do not sum up to the performance, the *α*-*β* tradeoff cannot be observed. (B) Comparison with UCM analysis. Three simulated strategies with no bias (i.e. with a mean action on the solution manifold) differ in performance because of differences in *β*, due to differences in the alignment of the covariance (*black* ellipses) with the Hessian (*red* degenerate ellipses), since the noise is equal). The decomposition of the action variability onto an *uncontrolled manifold* (UCM) subspace and an orthogonal (ORT) subspace does not discriminate the three strategies. (C) Comparison with GEM analysis. Three simulated strategies with equal noise but different performance due to differences in *α* are not discriminated by the *goal-relevant sensitivity* Σ_*GR*_ and the *goal-relevant variability fraction* Φ_*GR*_.

In contrast to both Hessian-based decomposition and TNC analysis, UCM analysis [27, 19] aims at decomposing action variability rather than characterizing the dependence of mean score on strategy and variability. It decomposes variability into a component within an *uncontrolled manifold* (UCM) subspace, which does not affect the task because such subspace is the null space of the Jacobian of the action-to-outcome function, and a component within the orthogonal (ORT) subspace. The ratio 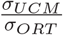, where *σ*_*UCM*_ is the square root of the variance in the UCM and *σ*_*ORT*_ in the ORT, then indicates whether a controller is successful in pushing variability onto task-irrelevant dimensions, thus improving performance. However, as shown by our decomposition, the relationship between action variability and performance, quantified by mean score, depends on the alignment of the action covariance with the Hessian and on the size of the Hessian rather than on the alignment with the Jacobian. Thus, a decomposition of variability onto the UCM and ORT subspaces does not discriminate strategies with different performance but equal *σ*_*UCM*_ and *σ*_*ORT*_ (and therefore equal ratio).

Fig 4B shows an example of three simulated strategies with different mean horizontal release velocity and different performances, due to different *β*, but equal *σ*_*UCM*_ and *σ*_*ORT*_. The three action distributions have means on the solution manifold, i.e. *α* = 0 and equal noise *η*. They also have the same orientation (with major and minor axes parallel to the 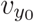 and 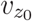 axes respectively) of the covariance ellipses but different ratios between the axes of the ellipses. As the Jacobian as well as the associated UCM and ORT directions change with the mean release velocity, the different amounts of variability along the horizontal and vertical velocities have the same amount of variability projected along the UCM and ORT directions. In contrast, the changes in the alignment of the covariance with the Hessian, which also changes with the mean release velocity, as well the changes in the size of the Hessian, do capture the features of the action distribution that affect performance. In particular, the differences in mean score, are due to different ratios of *θ* and *τ* as shown by the iso-lines in the *β* − *η* plane.

Finally, the GEM analysis [6] extends the UCM analysis by considering the noise sensitivity of the action-to-outcome function in addition to the directions of the Jacobian and its null space. The *total body-goal sensitivity* at the mean action, i.e., the ratio 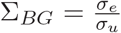, where *σ*_*e*_ is the square root of the total variation of the outcome 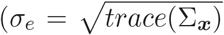 in our notation) and *σ*_*u*_ is the square root of the total variation of the action 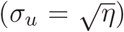, gives the amplification of variability between the action and outcome levels and can be decomposed as

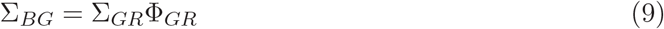

where the *goal-relevant sensitivity* 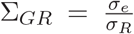, with *σ*_*R*_ = *σ*_*ORT*_, gives the amplification of variability in the ORT subspace and the *goal-relevant variability fraction* 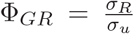 is the fraction of action variability available for mapping into the outcome and hence that can have an effect on performance. However, as for the UCM analysis, the GEM analysis characterizes the effect of the first order approximation of the action-to-outcome function, i.e., the Jacobian, on the variability in outcome space rather than the effect of the interplay between the geometry of the action-to-score function and the action distribution (both mean and covariance) on the mean score, i.e. the performance. Indeed, the GEM does not capture the effect on performance of a non-optimal mean in the distribution, which gives rise to the *α*-*β* trade-off presented above.

Fig 4C shows an example of three simulated strategies with the same horizontal release velocity and the same noise but different mean vertical release velocities and, thus, different *α* and different performances, as clearly shown in the *α*-*β* scatter plot. In contrast, the GEM decomposition does not discriminate between the three strategies, as they all have nearly equal Φ_*GR*_ and Σ_*GR*_. This is due to the fact that covariances are also nearly equal and the Jacobian does not change much when the mean vertical release velocity deviates from the optimal value on the solution manifold. However, such deviation of the mean action has a substantial effect on performance.

### 2.2 Unconstrained 6D throwing experimental data

To gain insights on individual strategies and their relationship to performance in a complex real-world motor skill we applied our Hessian-based decomposition to unconstrained throwing. We considered the release position and velocities collected from 20 participants performing several trials of unconstrained overarm throwing to four different targets (Fig. 1A) and the squared distance of the arrival position from the center of the target as the score (loss) of each trial. The performance, defined as mean score, varies significantly across participants (Fig. 1B, *p <* 0.001 linear mixed-effect model with participant and taregt identity as categorical predictors), and our aim was to characterize the key features of the individual action distributions that relate to performance and that differentiate participants with the same performance.

As actions are six-dimensional (6D), it is impossible to visualize both the score and the individual strategies in a single plot, as in the 2D throwing case. To illustrate some of the characteristics of the individual action distributions, Fig 5 shows the distribution of the three pairs of release velocity components in five exemplar participants (*columns*) aiming at target *T* 1. As there are 15 different pairs of 6 action variables (3 position and 3 velocity components) these plots do not fully illustrate the individual strategies (see Appendix B for a display of 9 of the 15 possible pairs). However, these examples show that the action distributions and the local geometry of the score function differ significantly across participants even when we consider a subset of action variables.

**Figure 5.**
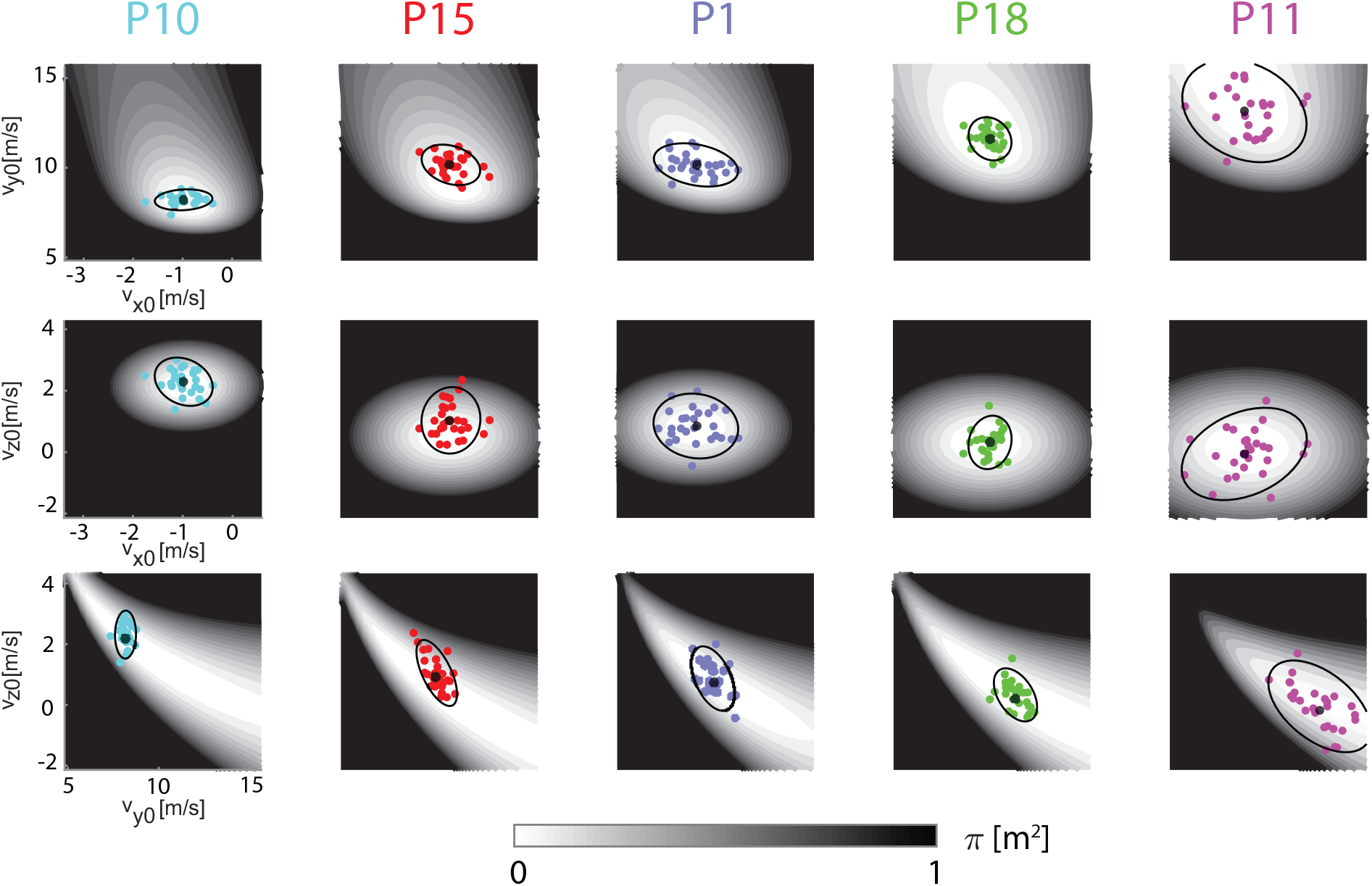
Examples of distribution of release velocity components for five representative participants (target T1). Each row illustrates a pairs of release velocity components 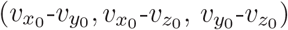. Colored circles represent release parameters of individual throws. Black circles and ellipses represent mean and covariance (95 % c.l.) of each parameter distribution. The gray-level shading of the contours indicates the local score, as a function of the release parameters. Note that in each row, the range of horizontal and vertical axes are the same for all participants, and the different shapes of the score across participant, reflect individual differences in the average action (mean position and velocity vectors at release). The wider the white area around the mean action, the more tolerant the score is to noise. Participants have been sorted from left to right according to their average release speed.

As for the simulated 2D strategies illustrated in Fig 3C, participants have been sorted from left to right according to their mean longitudinal release velocity 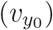, hence *P*10 is the slowest thrower and *P*11 the fastest. The domain of each velocity variable corresponds to the population mean ± 3 standard deviations. The score associated to each pair of velocity components, with the remaining velocity component and the three position components fixed at the subject-specific mean value, is indicated by the gray-scale shading of the contours. Note that the regions corresponding to actions with the same score (i.e. same gray shading) have different shapes (or geometry) across planes and participants, as they depend on the individual mean release action. The wider the white areas around the mean release parameter (*black large circle*), the more tolerant is the action-to-score function to action variability. In the 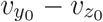 plane (*third row*), as in the 2D case, tolerance increases with mean 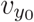. However, the mean longitudinal release velocity also affects the tolerance along the horizontal transverse release velocity 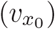, as can be observed in the 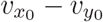 (*first row*) and 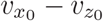 (*second row*) planes.

The distributions of the release parameters (*colored circles*), summarized in each plot of Fig 5 in terms of mean (*black large circle*) and covariance (two-standard deviations, *black ellipse*), also differs across planes and participants. In addition to differences in mean longitudinal velocity 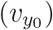, there are differences in mean vertical velocity. In particular, *P*15 (*second column*), on average, releases the ball with a faster vertical velocity 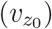 than *P*1, who has a similar longitudinal velocity. Remarkably, individual covariances differ both in size and orientation. In the 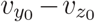 (*third row*) plane, *P*10 shows a small covariance ellipse with the major axis oriented along the 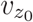 axis. In contrast, *P*11 shows a much larger covariance ellipse with the major axis oriented diagonally, i.e. with a negative correlation between the two velocity components.

In sum, in all three pairs of velocity variables (as well as in the other pairs of variable shown in Appendix B), both the local geometry of the action-to-score function and the structure of action variability suggest that specific features of the individual action distributions affect performance, as they determine the amount of overlap of the distribution with the regions with lowest penalty. However, as the distribution is 6-dimensional and there are 15 different pairs of action variables, it is not possible to identify by visual inspection a unique source of the inter-individual differences in throwing strategies and to systematically explain the relationship between action distribution and throwing performance. These limitations can be overcome by the Hessian-based decomposition.

#### Validity of the Hessian-based decomposition

The Hessian-based performance analysis is based on the assumption that the individual action distribution is sufficiently localized around the mean action such that higher order terms of the Taylor expansion can be neglected in the action score approximation. To validate this assumption with our experimental data, we tested whether Eq (8) can be considered as an acceptable model of the mean action score. Fig 6 shows the relationship between the mean squared error and the sum *α* + *β* across participants and targets. We found that the sum *α* + *β* could explain 99% of the variance for targets T2, T3, and T4 and 97% for target T1, due to the larger error observed in *P*9. *P*9 had in fact the worst performance (see Fig 1B) and the largest bias (*α*) and variability (*η*), which violates the assumption of the action distribution being sufficiently localized. In sum, *α* + *β* can explain quite accurately the individual performance across participants and conditions, except for *P*9, where the sum *α* + *β* tends to overestimate the average score for T1,T2 and T3.

**Figure 6.**
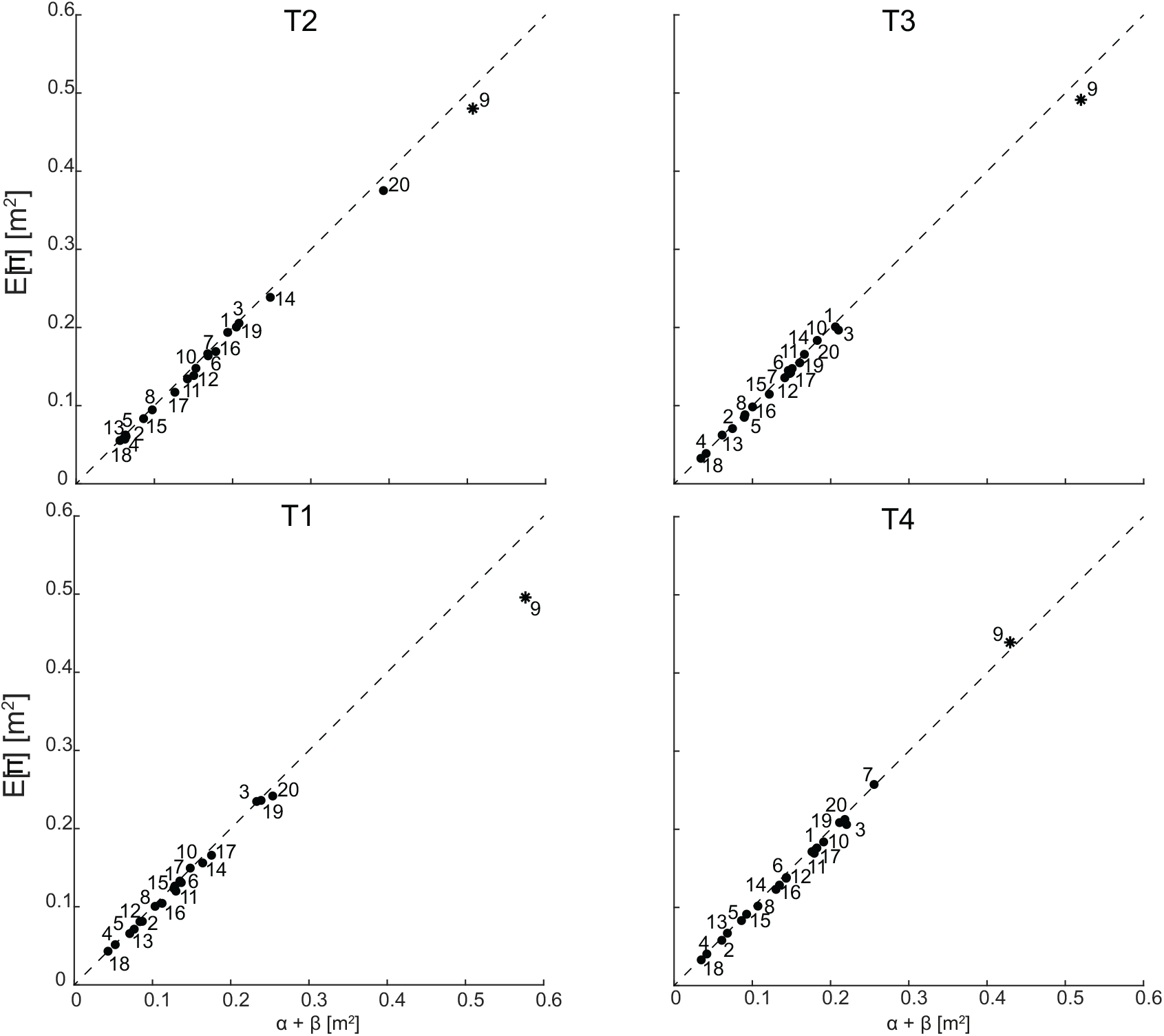
Validity of the quadratic approximation. Mean score vs. quadratic approximation (8) across targets (T1-T4, *different panels*) and participants (*P*1-*P*20) as in 1. Note that *P*9 is the only participant deviating substantially from the identity line (*dashed line*).

#### Hessian-based decomposition of the mean score reveals key performance-related features of individual throwing strategies

The Hessian-based decomposition of the mean score relies on the straightforward computation of the action mean, action covariance and Hessian of the action-to-score function at the mean action. While the analysis of the structure of the Hessian and the action covariance matrices provides useful information on the task geometry and on the features of the action variability shared across subjects (see Appendix C), the parameters extracted from the Hessian-based decomposition of the mean score of Eq (8) compactly describe the key performance-related features of the individual action distribution and allow to fully characterize how performance depends on such features.

Fig 7 shows the distributions of all the parameters of the Hessian-based decomposition of the mean score (*α, β, η, τ* and *θ* defined in Eq 24) across participants and relative to target T1. To visualize the inter-individual differences in terms of these five parameters, as for 2D throwing, we consider four scatter plots. The five exemplar participants of Fig. 5 are identified by the color of the edges of the circular markers and show in 6D throwing strategies that are similar to those illustrated in 2D throwing examples of Fig 3, with matching colors.

**Figure 7.**
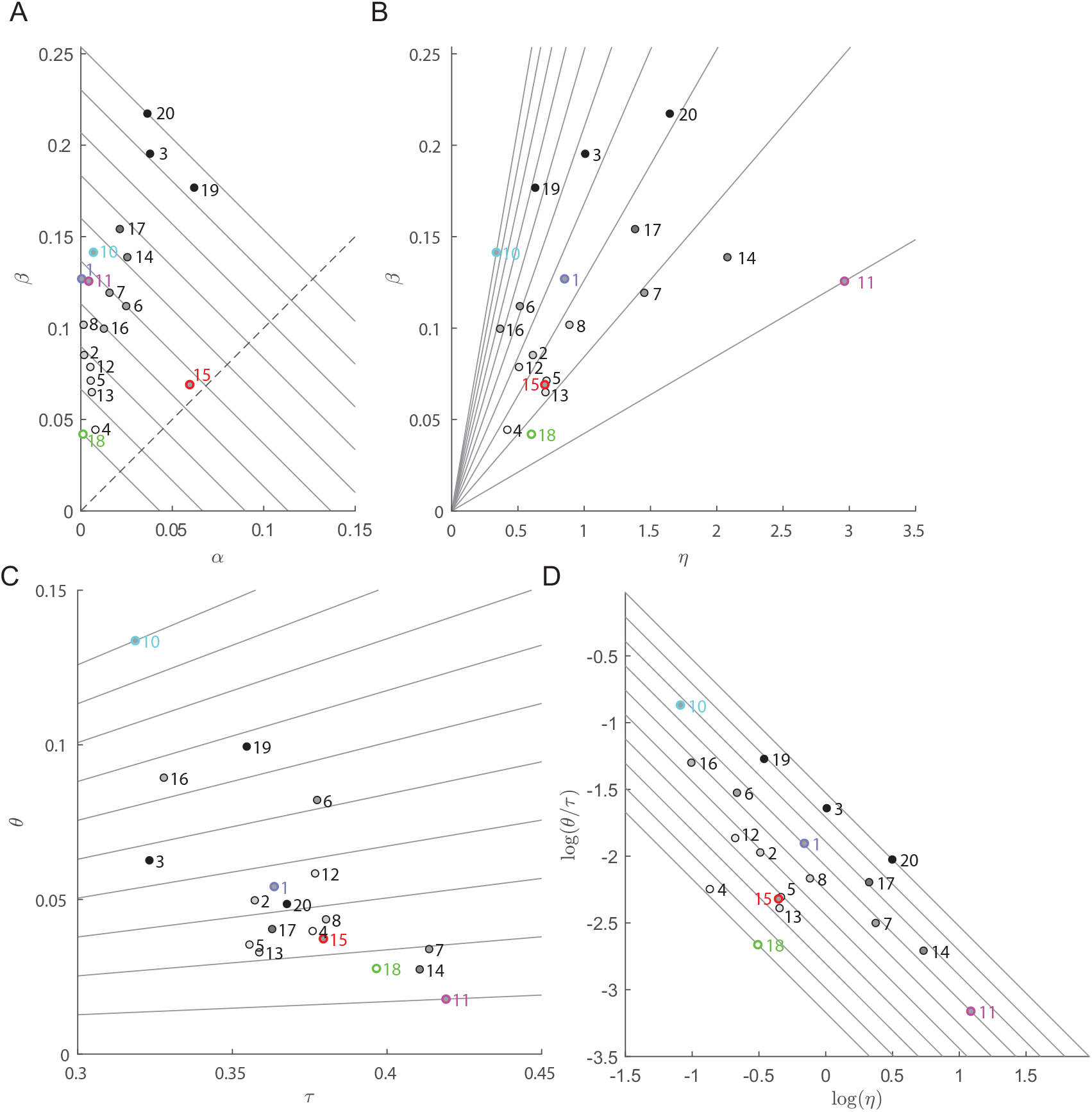
Decomposition of the mean score across participants for target T1. Circular markers are filled with areas that are gray-shaded according to the score (lighter gray for lower mean score). The edge of the markers corresponding to the five participants of Fig 5 are colored with matching colors. To increase the readability of the figure, *P*9 is not shown. (A) The *α* - *β* plane and the iso-performance lines. The dashed line represent the direction of maximal change in performance (orthogonal to the iso-performance lines). (B) The *β* - *η* plane. The lines have slopes corresponding to different values of 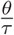. Participants along the same line, such as *P*1, *P*2, and *P*12, have the similar alignment-to-tolerance ratio and the difference in their *β* are only due to the uncorrelated noise *η*. (C) The *θ* - *τ* (tolerance-alignment) plane. The lines have slopes corresponding to different values of 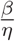. The negative correlation between *θ* and *τ* indicates that participants with mean release action in a region with higher tolerance also tend to have lower alignment with the most noise sensitive directions. (D) The *log*(*η*)-*log*(*θ/τ*) plane and iso-*β* lines. As *β* is the product of *θ/τ* and *η* and *log*(*β*) = *log*(*θ/τ*) + *log*(*η*), in the *log*(*η*)-*log*(*θ/τ*) plane participants with the same *β* lie on a line with a -1 slope and maximal change of the tolerance-variability index occurs along the orthogonal direction.

The *α* − *β* plane (Fig 7A) characterizes the performance of each participant and the trade-off between bias and tolerance-variability. The *diagonal continuous* lines with slope −1, corresponding to equal sum *α* + *β*, indicate iso-performing strategies. Participants closer to the origin of the plane are the best performers (e.g., *P*18 and *P*4, compare with Fig 1B). Moving away from the origin in the direction of the *dashed* line performance decreases. Participants with the same performance, i.e. on the same diagonal line, differ on their position on the line, thus showing a different trade-off between bias (*α*) and tolerance-variability (*β*). For example, *P*10, *P*1, and *P*11 have low *α* and *P*15 high *α* but they are all close to the same iso-performance line and thus have similar performance. In sum, the *α* − *β* plane shows that our participants differ in their performance because of their specific values of the *α* and *β* indicators and that participants with the same performance trade-off *α* with *β*.

The *η*-*β* plane (Fig 7B) characterizes the role of the amount of action variability, i.e. the uncorrelated noise *η*, in determining the tolerance-variability index *β*. The same contribution to performance due to tolerance-variability may be obtained with very different amounts of variability. For example, *P*10, *P*1, and *P*11 have similar *β* values but very different *η* values. This is because larger variability is compensated by lower alignment and higher tolerance. In fact, the slope of the lines in the *η*-*β* plane corresponds to 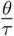 and participants on lines with lower slope have a lower alignment or higher tolerance.

The interplay between tolerance and alignment is illustrated in the *τ* -*θ* plane (Fig in 7C). Here the slope of the lines corresponds to a constant 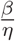, i.e. each line corresponds to a line in the *η*-*β* plane (as 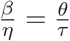). While the relationship between *τ* and *θ* is not fixed, participants with a higher tolerance also have a lower alignment. This indicates that, when the mean action is located in a more tolerant region of the action space (e.g. higher horizontal velocity, as for *P*11) it is also possible to further optimize performance by reducing the alignment.

Finally, the *log*(*η*)-*log*(*θ/τ*) plane (Fig 7D) characterizes the trade-off between the amount of action variability and the other features of the action distribution determining the tolerance-variability index. In the log-log plane the multiplicative contribution of 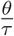 (determined by the mean action and the alignment of the action covariance) and of the amount of action variability *η* to *β* are represented by a sum 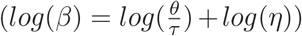 and, thus, strategies with equal *β* are located on lines with slope −1. Then, this plot clearly shows how participants differs in their tolerance-variability index *β* (participants with lower *β* such as *P*18 and *P*15 are on lines in the *bottom-left*) and how participants with the same *β* trade-off noise with tolerance and alignment (e.g. *P*10 vs. *P*1 vs. *P*11).

#### Consistency of the decomposition of individual action strategies across targets

To test whether the individual performance-related features of action distributions described by the Hessian-based decomposition are consistent, we compared the *α, β, η, τ* and *θ* parameters across both participants and targets. If these parameters are robust indicators of the inter-individual differences in throwing strategy, we expect them to vary significantly across participants but not across targets.

Fig 8 shows the distributions across participants and targets of the parameters of the Hessian-based decomposition in the same combinations of parameters as in Fig 7, which included only target T1. The quadrilaterals (*dashed edge lines* and *color-shaded areas*), with the parameters for the four different targets of each participant as vertexes, are colored according to the mean score rank of each subject. In most cases the parameters do not show large variations across targets, i.e. they represent consistent features that characterize individual strategies. In the *α*-*β* plane (Fig 8A), a few participants show large across-target variations (e.g. *P*14, who has a larger *α* for T2 than for the other targets; *P*7, who has a larger *β* for T4; *P*20, who has both *α* and *β* larger for T2). However, most participants occupy small and distinct regions of the *η*-*β* (Fig 8B) and *log*(*η*)-*log*(*θ/τ*) (Fig 8D) planes. Finally, in the *τ* -*θ* plane (Fig 8C) about half of the participants occupy the region with intermediate values of *τ*.

**Figure 8.**
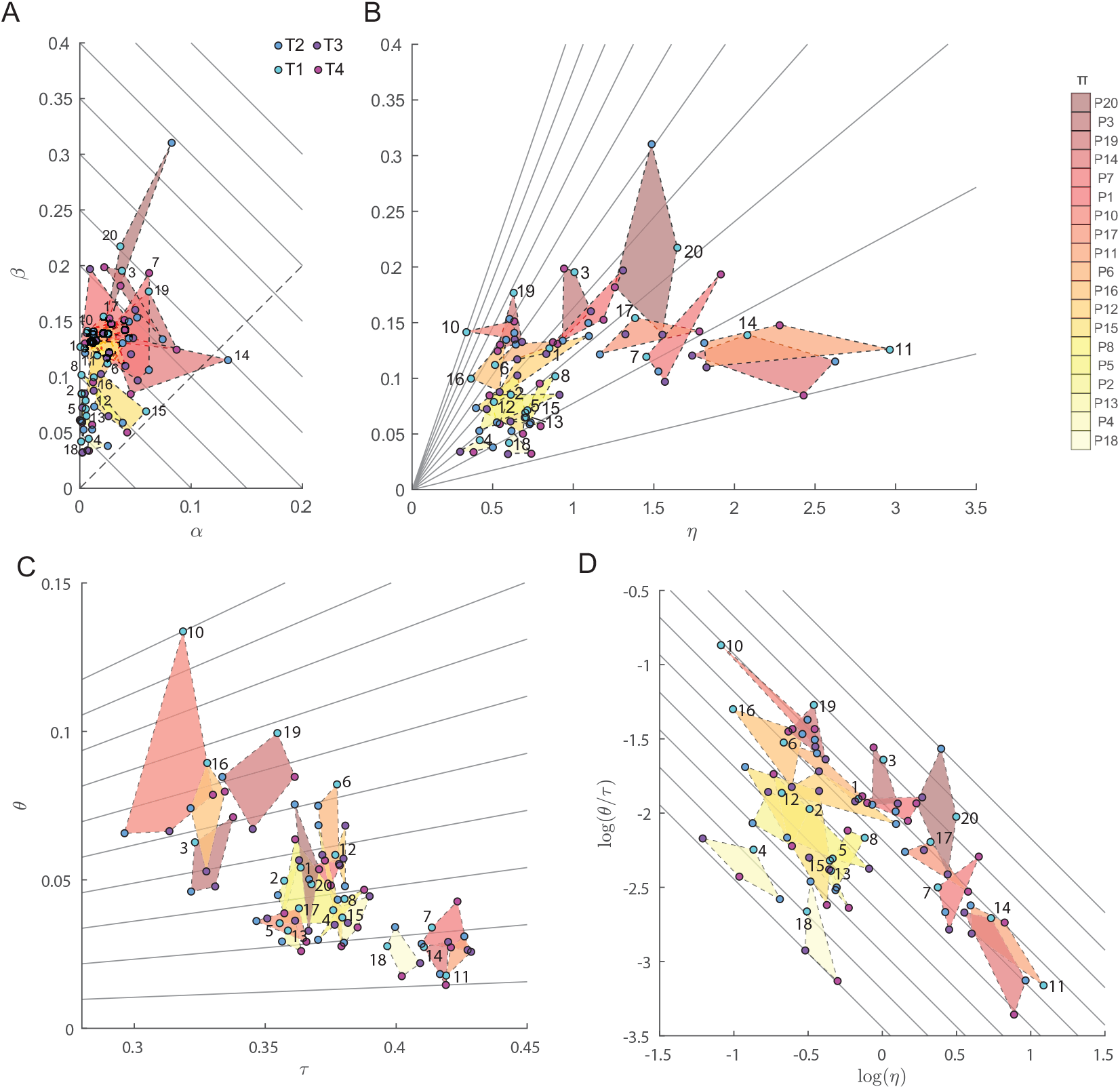
Decomposition of the mean score across participants for all targets. In each panel, the quadrilateral connects the points corresponding to the values of a pair of indicators for the four targets of each participant (excluding *P*9) and they are colored according to the mean score of that participant (color scale on the *top right*). The targets are indicated by different colors of the circular markers and the participant number is indicated close to the marker for T1 (except for *P*7 and *P*14). (A) The *α* - *β* plane and the iso-performance lines. (B) The *β* - *η* plane. (C) The *θ* - *τ* (tolerance-alignment) plane. (D) The *log*(*θ/τ*)-*log*(*η*) plane.

A linear mixed-effect model with with participant and target identities as categorical predictor (Table 1) of all five parameters supports these qualitative observations, showing that the effect of participant is significant for all parameters while the effect of target is significant only for *α* (*P*= 0.02) and *τ*. Thus, *α* and *τ* are the only parameters that are not consistent across targets. This is not surprising for *τ* since it depends on the individual strategy only through the mean release position and velocity, which, in turn, depend on the target position, and, thus, *τ* varies with the target. Concerning *α*, the contrast between pairs of targets (fixed effect coefficients) shows that its value is significantly higher for *T* 2 than *T* 1 (Δ*α* = −0.02, *p <* 10^−2^) and *T* 3 (Δ*α* = −0.01, *P*= 0.01), indicating that the bias of a throwing strategies also depends slightly on the target.

**Table 1.**
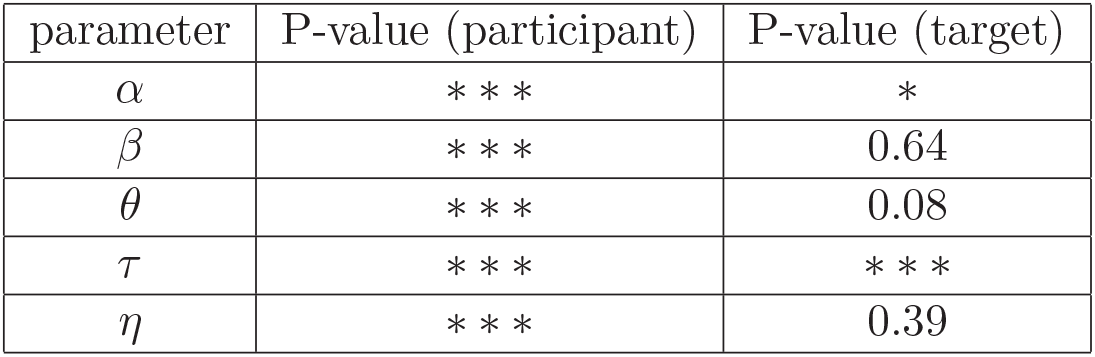
Robustness of decomposition parameters across targets. P values of a linear mixed-effect model with with participant and target as categorical predictors. *: *p <* 0.05; **: *p <* 0.01; ***: *p <* 0.001.

| parameter | P-value (participant) | P-value (target) |
| --- | --- | --- |
| $\alpha$ | *** | * |
| $\beta$ | *** | 0.64 |
| $\theta$ | *** | 0.08 |
| $\tau$ | *** | *** |
| $\eta$ | *** | 0.39 |

## 3 Discussion

We have developed a novel method to investigate how the distribution of actions in goal directed behaviors relates to individual performance. The method allows to characterize how performance depends on a few critical features of the action distribution, for tasks in which actions are redundant (the same goal may be achieved by multiple actions), high-dimensional (each action is described by a vector with many components) and noisy (actions vary due to stochastic sensory and motor processes). Assuming that the success of an action can be assessed by a scalar cost, i.e. a score, and that such score is a smooth function of the action, we derived an approximate analytical relationship between the mean score and the first two moments of the actions distribution: the mean action ***ā*** and action covariance Σ^*a*^ across multiple trials. Performance, defined as the mean score, can be approximated as the sum of two components: the score of the mean action (*α*(***ā***)) and a tolerance-variability index (*β*(***ā***, Σ^*a*^)). The *α* parameter, when different from zero, measures deviations of the mean action from the set of actions that accurately achieve the goal (solution manifold). The *β* index, instead, measures how the mean score is affected by actions variability and by the geometry of the action-to-score function (determining the sensitivity to noise as a result of the non-linearities of the action-to-score function around the mean action and their alignment with the directions of largest variability in the action distribution). Such index results from the product of three terms: (i) the *uncorrelated noise* (*η*), computed as the sum of the variances of the individual components of the action vector (i.e. the trace of the action covariance matrix); (ii) the *tolerance* of the score function (*τ*), quantifying the overall sensitivity of the score to deviations from the mean action due to the curvature of the action-to-score function captured by the Hessian; (iii) the *alignment* (*θ*), a scalar measure of the effect on mean score of the relative orientation between the most sensitive directions of the action-to-score function and the directions of maximum variability of the actions. Thus, these five indicators provide a compact yet informative characterization of the features of the action distributions that affect performance in relation to a specific task, and allow to capture detailed facets of individual strategies in goal directed behaviors.

We have applied this method to characterize individual performance and variability in unconstrained overarm throwing actions of twenty non-trained participants. Across participants there were remarkable inter-individual differences in the *α, β, η, τ* and *θ* parameters (Fig 7) and most of these differences were consistent across targets (Fig 8). In line with previous works focusing on low-dimensional throwing tasks [23, 6], in our unconstrained high-dimensional throwing task we found that skilled participants have small *α* (accurate mean action) and small *β* (tolerance-variability index). Still, it is possible to further differentiate different optimizing strategies, as low *β* can be achieved by minimizing action variability or by compensating the higher variability in action execution (*η*) with higher tolerance *τ* and smaller alignment (i.e., smaller *θ*). Moreover, examining the scatter plot of different pairs of parameters of individual participants, we identified specific combinations of parameters that do not affect performance, but correspond to different throwing strategies. First, similar performance can be achieved trading off bias (*α*) with tolerance-variability (*β*). Second, similar *β* can be achieved trading off alignment and tolerance (*θ/τ*) with variability (*η*). Such interplay between variability and geometric features of the score can be observed in several pairs of action variables (e.g. in the *v*_*y*_-*v*_*z*_ plane, Fig 5) but are characterized, fully yet compactly, by the indicators of the decomposition (Fig. 7).

### 3.1 Comparison with related approaches

In redundant, high-dimensional, and noisy tasks it is not enough to characterize the mean and the covariance of the action distribution to fully capture the relationship between action variability and performance. In agreement with earlier computational approaches addressing variability in multivariate actions [27, 19, 23, 15], our method highlights the key role of the geometry of the action-to-score function (captured by the Hessian) to assess how action variability affects performance. Differently from first-order methods such as UCM, GEM, or the more recent approach in [34], which characterize the local geometry with a linear approximation (expressed through the Jacobian matrix or the gradient vector), our method relies on a second-order approximation (based on the Hessian matrix). The main reason for which our method does not depend on the first-order term of the Taylor expansion of the action-to-score function and requires a second-order approximation is the fact that we are considering the mean score rather than the variability in action or outcome space as a measure of performance. As indicated in equations (14) and (15), the mean score does not depend on the gradient of the action-to-score function computed at the mean action, the reason being that the first-order term of the expansion is multiplied by the mean deviation from the mean action, which is null by definition. In other terms, changes in score (with respect to the mean) associated to actions that deviate from the mean action sum up to zero in the linear approximation of the action-to-score function. Indeed, for a linear action-to-score mapping the mean score is given simply by the score of the mean action, as all higher order derivatives in the expansion are null. Thus, in a quadratic approximation, it is only the local curvature of the action-to-score function, captured by the Hessian matrix, that affects the mean score.

How action distribution affects the mean score in a goal-directed behavior has been addressed by the TNC methods [23, 5, 31, 30]. The methods have been developed for, and applied to, a two-dimensional throwing task inspired by the skittle game, in which participants have to hit a target by releasing through a rotating joint (i.e., the action parameters are the release angle and tangential velocity) a virtual ball that could rotate in a plane around a pole. The TNC methods take into account the geometry of the action-to-score mapping implicitly, by evaluating the effects on performance of different action distributions through surrogate data (when comparing pairs of distributions [23]) or by exhaustive search (when comparing with optimal distributions [5]). In contrast, our method explicitly decomposes the contribution of different features of the action distribution through a Taylor expansion. Such analytical approach overcomes the disadvantage of the TNC methods concerning the use of numerical procedures necessary to generate surrogate data or to search for optimal distributions, which limits its applicability to high-dimensional actions. Moreover, our method allows to determine the contribution of the local geometry and of the action variability independently on each other. As illustrated in the 2D throwing examples of Fig 4A and detailed in the Appendix (see S1 Appendix A), under the assumption of a smooth action-to-score function, for which the Hessian matrix is well defined, the tolerance, noise, and covariation terms of the TNC decomposition correspond to specific combinations of the terms in our decomposition. However, importantly, all the three terms of the TNC decomposition depend on both the Hessian and the action covariance, thus they do not indepentently characterize the contributions of the local action-to-score geometry, of the total action variability, and of the alignment between action variability and score tolerance *τ, η*, and *θ*. Moreover, when assessing a single distribution, the TNC-Cost analysis does not provide a unique decomposition of performance, as the three costs do not sum up to the mean score. Then, it is not possible to directly characterize the trade-off between the different components of the variability and local task geometry as with the Hessian-based decomposition. As an advantage, however, the TNC method does not rely on any assumption on the action-to-score function, such as smoothness and adequateness of a second-order approximation.

A fundamental difference between GEM and both the TNC and our approach, is the choice of performance measure. In the study of goal-directed motor skills performance is typically associated with a measure of *error* with respect to a predefined goal. Reduction of task errors and their variability is broadly recognized as an indicator of skilled performance. However, more recent views of human motor control, based on decision-making theory, propose an alternative definition of performance which is based on the concept of a score/loss function that essentially assigns a number (or a cost) to a given task error. In this perspective, the *mean* or *expected loss* is taken as measure of performance over repeated trials of a given motor task. Central to the GEM analysis is a goal function *e*(***a***) = ***x***_*T*_ − ***f*** (***a***) that expresses the error between a desired goal/target *x*_*T*_ and the outcome *f* (***a***) of a given action *a*. By linearizing this error/goal function around the mean action of a strategy, GEM quantifies the overall contribution of *tolerance, noise* and *alignment* with the task/goal-level variability, i.e *var*(*e*). The Hessian-based decomposition proposed in this work extends the GEM analysis to decomposition of the mean of a performance indicator. In particular, we have shown that is the Hessian and not the Jacobian that affects the expected score/loss around the mean action of a strategy (see equation (30)).

### 3.2 Assumptions and limitations

Our decomposition method requires a smooth action-to-score function and it provides an accurate estimate of the mean score only if the non-linearities in such function are adequately approximated by the second-order term of the Taylor expansion over the domain spanned by the actions. The assumption of smoothness (or at least continuity of the function and all partial derivatives up to the second order) is valid for a broad class of score functions, such as most penalty or reward functions usually employed to quantify task performance. For goal-directed tasks, requiring to minimize the distance from a target, the squared distance is a good choice because it leads to an action-to-score function which is twice differentiable everywhere the action-to-outcome function is smooth. The squared distance is preferable over the Euclidean distance because the latter has a singularity in the second derivative at zero, i.e. on the solution manifold. However, if the subject is attempting to minimize (maximize) the score, as the distance and the squared distance have the same minimum (maximum), both functions capture the control strategy equally well.

Another key assumption in our approach is that the second-order Taylor expansion of the action-to-score function around the mean action provides an acceptable approximation. As shown in Fig 6, for almost all participants and targets, the estimation of the mean score based on such approximation (*α* + *β*) is close to the actual mean score (*E*[*π*]). The only exception is participant *P*9 who had a poor performance and a very large variability in the ball release parameters. Indeed, the validity of the quadratic approximation depends on the nature of the non-linearities of the action-to-score function and the range of the deviations from the mean, i.e. from the relative spatial scales characterizing the concentration of the action distribution and the Hessian. Thus, if behavior is very erratic, our decomposition may become inaccurate for the entire set of actions and may be restricted to a more concentrated subset. However, considering that participants in our sample were untrained throwers, it is noticeable that the quadratic approximation was good for all but one out of twenty participants. This suggests that our methods could be safely applied to more controlled tasks, e.g. in evaluating athletes performances (as athletes do not typically exhibit high variability in motor actions) or in assessing motor skill learning, where training tends to quickly reduce motor variability. However, in cases where the variability is high, such as during the initial exploration phase when learning a novel motor skill, one could apply the decomposition on local clusters of data and still provide a compact characterization through the parameters of Hessian-based decomposition of each cluster. We plan to develop such approach based on a mixture of Hessian-based decompositions and to apply it to the investigation of throwing skills learning in future work.

Our decomposition method relies on the computation of the action covariance and the Hessian of the action-to-score function. These matrices and some of the parameters of the decomposition depend on the choice of the coordinate system in action space. In particular, the noise *η* and the tolerance *τ*, being defined as traces of the covariance and Hessian matrices, respectively, change under coordinate transformations (unless a metric is chosen [3]). However, the *α* term is a scalar (i.e. is a single number corresponding to the score associated to the mean action) and it does not depend on coordinates. The *β* term is the trace of the product of the covariance and the Hessian matrices and it is invariant under affine coordinate transformations, given that Σ^*a*^ and *H*^*a*^ transform in opposite ways (see S1 Appendix B). Thus, re-scaling of positional and velocity coordinates due to different choices of measurement units does not affect the decomposition of mean score as a sum of *α* and *β*. However, *β* is not invariant, in general, for non-linear coordinate transformations, such as the transformation from Cartesian to polar coordinates. Indeed, the dependence on action coordinates has raised concerns about the reliability of the TNC decomposition [29, 1] and of UCM and GEM methods [32]. While such dependence may provide an opportunity to evaluate the role of different coordinate systems for control [22], it has also been noticed that geometric properties of the action-to-score function such as the solution manifold do not depend on coordinates [32]. In our decomposition, if the mean of the action distribution is on the solution manifold (*α* = 0), *β* is invariant also under non-linear transformations, because the non-linear term in the transformation of *H*^*a*^ depends on the gradient of the action-to-score function, which is null on the solution manifold. Moreover, if the action distribution is not centered on the solution manifold but it is concentrated (i.e. *η* is small) the change in *β* due to non-linear coordinate transformations may be negligible.

A further limitation of our approach, which is shared with GEM and TNC, is that the decomposition cannot reveal the temporal structure of inter-trial fluctuations, as multiple trials are needed to compute the mean and the variance of an individual’s action strategy. Recent work addresses this issue with an inter-trial error correction model that predicts both the temporal and geometric structure of variability near the goal equivalent manifold (GEM) of a simplified shuffle board task [16]. Furthermore, the variability analysis is shown to be coordinate-independent as the characterization is performed in the eigenspace of the error-correcting controller matrix.

### 3.3 Applications to motor skill learning

In this work we have focused on characterizing steady-state performance and individual action distribution during short experimental sessions rather than on skill improvement over multiple sessions. Future work will include longitudinal studies to understand if and how the observed inter-individual differences are related to the time course and the magnitude of individual performance improvements and skill learning. Current theories of human sensorimotor control suggest the existence of two distinct mechanisms underlying motor skill learning: a model-based system that improves motor performance guided by an internal forward model of the body and the environment, which is updated based on prediction errors [28]; and a model-free system in which learning is driven by reinforcement and punishment of successful/erroneous actions [11, 4]. Motor adaptation studies, in which a systematic perturbation of the environment is introduced by means of force fields of visuomotor rotations, suggest that the model-based system is responsible for the quick adaptation/compensation of the mean error. The model-free system, driven by reinforcement and punishment, regulates instead motor variability, and is hence responsible for the slow reduction of the variable errors. However, the interplay between this two learning mechanisms, remains poorly understood.

In analyzing our free overam throwing data, we have highlighted the existence of iso-performing participants, such as *P*1, *P*11 and *P*15, which have the same mean score, but different contributions of *α* and *β*. Do inter-individual differences in terms of *α* and *β* translate into individual differences in terms of performance improvement? In future work we plan to use the proposed framework to study the acquisition of throwing skills in virtual reality environments in which we can alter both the dynamics of the ball, for instance by manipulating the (virtual) gravity field, as well as the task score geometry, in this work assumed quadratic and isotropic in both task directions. As adapting to an altered dynamics requires learning a new forward model while a new task geometry changes the reward function, the dissociation between these two contributions might allow us to dissociate between model-based and model-free learning and to understand how initial inter-individual differences in terms of performance, variability and score tolerance translate into individual performance improvement.

### 3.4 Summary and conclusions

We have introduced a novel method to characterize the key features of the distribution of high-dimensional and redundant actions that affect performance, defined as the mean of the score assigned at individual actions. We have applied the method to the investigation of inter-individual differences in unconstrained throwing. We found that the indicators derived from the Hessian-based decomposition allow to identify specific and consistent features relating individual throwing strategies to different performance level and to understand how different strategies achieve similar performances. Participants differed in their throwing performance because they consistently differed in either the score of their mean action (bias, *α*) or the level of their tolerance-variability index (*β*), a measure of the interplay between action variability and action-to-score geometry. The same performance could be achieved trading off *α* with *β* and the same *β* could be achieved trading off *θ/τ* with *η*. In sum, the compact characterization of the relation between high-dimensional, redundant, and noisy action distributions and performance provided by our Hessian-based decomposition may be applied to a variety of complex real-life motor skill, opening up new opportunities both for systematic investigations of inter-individual differences in real-life motor skills and for practical applications to training of complex motor tasks. As motor control investigators, we plan to address in future studies how the individual parameters of the Hessian-based decomposition relate to individual motor learning capabilities, a first step to unravel the fundamental mechanisms underlying individual learning differences. A sport trainer could also use the Hessian-based decomposition indicators of a trainee, based on the analysis of a few tenths of repetitions of a specific motor skill, to select an optimal individualized training strategy according to those indicators.

## 4 Methods

### 4.1 Performance as expected value of non-linear and high-dimensional action scores

Let us assume that, at every trial *t*, an individual generates an action *a*_*t*_ ∈ ***A*** ⊂ *R*^*n*^ with some random noise such that the action distribution can be described by a probability density function (p.d.f.) *p*_***A***_. Let us also assume that, at every trial, the action receives a score point *π*_*t*_ = *s*^*a*^(***a***_*t*_) through a *score* (or loss) function *s*^*a*^ : ***a*** ∈ *R*^*n*^ → *R*^+^ which punish motor actions according to some optimality criteria such as task errors or metabolic cost of an action. For instance, in Fig 1, the score function assigns a penalty score to an action, that is the squared distance between the action outcome ***x***(***a***) and the target position ***x***_*T*_. We define the *performance* of a motor strategy *p*_***A***_ as the *expected score*:

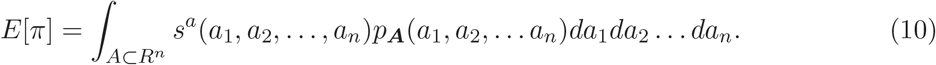

which is the sum over all possible actions of the probability of a taking a given action times its score (integrated in *R*^*n*^). In this framework, an individual motor strategy *p*_***A***_ is optimal if, over repeated attempts, minimizes the *average* score. Similarly, given two motor strategies we can establish if they are *equivalent* or if one is better than the other by simply comparing their expected score. In real practice, the score function may be highly non-linear and the action space high dimensional (*n >>* 1) making difficult to find an analytic solution to (10). Therefore, we do not usually solve (10) but instead approximate the expected score with its sample average as in Fig 1B, where best performers, such as *P*18 and *P*4 have the lowest average quadratic error. Nevertheless, the sample average by itself is not informative enough to understand *why* the individual action distributions of *P*18 and *P*4 are much more performing compared to the majority of participants.

#### Quadratic approximation of the expected action score

By restricting the action score to the class of continuous and at least twice differentiable functions, and assuming that the action distribution is sufficiently localized around the mean action, it is possible to find an approximate but analytic solution to (10).

Let’s assume that an individual selects motor actions according to a p.d.f. with *expected* or *mean action E*[***a***] = ***ā*** ∈ *R*^*n*^, and covariance matrix Σ^*a*^ ∈ *P* ^*n*^ (symmetric and positive definite). In other words, at every trial *t*, an individual generates an action *a*_*t*_ according to the following stochastic model:

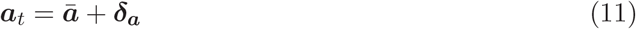

where ***δ***_*a*_ ∈ *R*^*n*^ is the *stochastic component* of the individual strategy, which, at every trial, ‘perturbs’ the mean action ***ā***. The mean action represents an individual preference in choosing, *on average*, a given action and the covariance matrix Σ^*a*^ = *E*[***δ***_*a*_^*T*^ ***δ***_*a*_] represents action covariation/correlation (*action variability*) across multiple trials.

Let us also assume, that locally, i.e. in a neighborhood of the average action ***ā***, the score *s*^*a*^(***a***_*t*_) of an action ***a***_*t*_, can be approximated with the following second-order Taylor expansions:

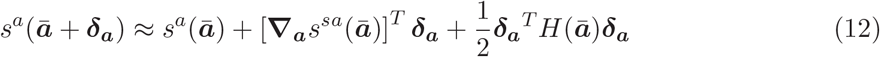

where 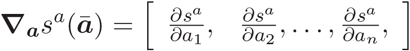 is the gradient of the action score function evaluated at the average action ***ā*** and:

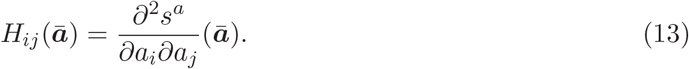

is the *n* × *n* Hessian matrix (assumed to be symmetric and positive definite) of the action score evaluated at ***ā***.

Inserting (12) in the integral (10), we can write the expected action score as the sum of three terms:

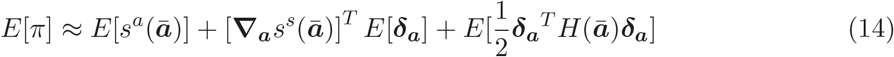

the first term *E*[*s*^*a*^(***ā***)] is simply *the score of the average action s*^*a*^(***ā***) given that the expected value of a constant is the constant itself. The second term, [**∇**_*a*_*s*^*s*^ (***ā***)]^*T*^ *E*[***δ***_***a***_], vanishes whenever the quadratic approximation is evaluated at the mean action ***ā***, given that in such condition *E*[***δ***_*a*_] = 0. The last term corresponds to the expected value of a *quadratic form*, which is well known to be equal to 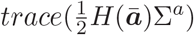 [12]. This term is zero: i) when the score is a linear function of the action, in which case the Hessian is zero, ii) when actions are not stochastic Σ^*a*^ = 0, or when *H* and Σ^*a*^ are ‘orthogonal’. In all other cases this term will influence the expected action score.

Under such hypothesis the expected action score (10) can be (locally) approximated as:

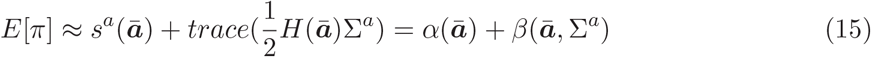

where:

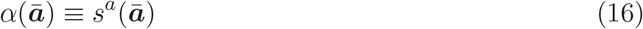

is *the score of the average action*, and

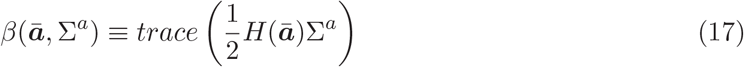

is the *tolerance-variability index* which captures how local non-linearities of the action score (*H*(***ā***)) and variability in the action strategy (Σ^*a*^) affect (increase/decrease) performance, i.e. the mean score *E*[*π*]. Note that the definition of *β* corresponds to half the Frobenius inner product between the Hessian and the Covariance matrix. In the next section we show that it can be decomposed into three independent components: *the uncorrelated noise, the score tolerance* and the *alignment*.

### 4.2 Hessian-based decomposition of motor skills

#### Principal variability directions and *uncorrelated noise η*

The action covariance matrix Σ^*a*^, symmetric and positive-definite, can be decomposed (principal component analysis) into singular values (or eigenvalues) and singular vectors (or eigenvectors):

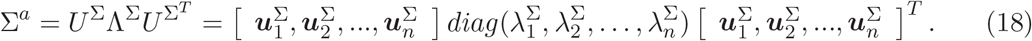

with 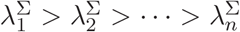. The larger a singular value 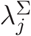, the more variable will be the action strategy along its associated *principal variability direction* 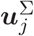.

We define the *uncorrelated noise η* of an individual strategy as the total variation of the action distribution:

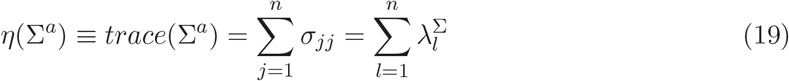

#### Principal sensitivity directions and *tolerance τ*

When locally, i.e. around the mean action ***ā***, the score is a continuous and at least twice differentiable function of the action variables, the Hessian is an *n* × *n* symmetric matrix (Schwarz’s theorem) which can be written as:

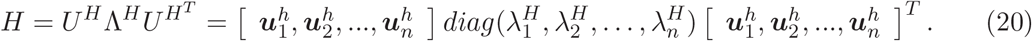

The diagonal matrix 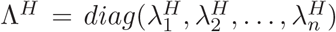 contains the *n* singular values 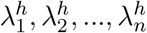 (with 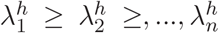) of the Hessian matrix. The orthonormal matrix *U*^*H*^ contains the associated singular vectors 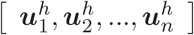. The larger a singular value 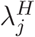, the greater will be the change of the score (i.e. the greater the curvature will be) along its associated *principal sensitivity direction* 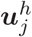. When the Hessian matrix is positive-definite, i.e the singular values 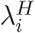 are positive, the score function is locally convex and therefore the mean action is in, or ‘close to’, a (local/global) minimum of the score function. Conversely, negative eigenvalues are representative of concave regions of the score function, while eigenvalues with mixed signs suggest that the average action is in/close-to a saddle point of the score function. In this work we will focus on score functions which are locally convex and for which the Hessian matrix is semi positive-definite, i.e. all eigenvalues are greater or equal than zero, although what follows can be generalized to more complex score functions having a landscape with many minima, maxima and saddle points.

The eigenvalues 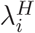 express the *sensitivity* of the score to stochastic perturbations. The larger 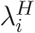, the more sensitive the score function is to perturbations ***δ***_*a*_ which are directed along the *i*-th principal sensitivity vector 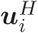. As a local, scalar measure, of ‘total curvature’ of the score, we define the *sensitivity* of the score as the 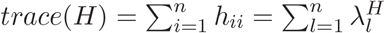.

The local *score tolerance* then, is defined as the reciprocal of the score sensitivity:

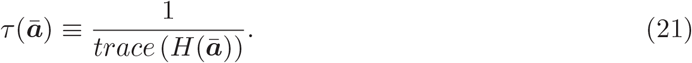

##### Alignment θ

With the above definitions of tolerance and noise, by normalizing both the Hessian and the covariance matrix by their respective traces, we can rewrite the tolerance-variability index *β* as:

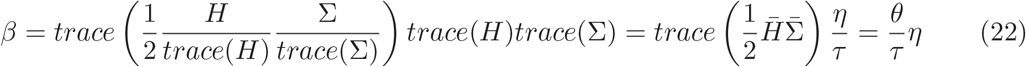

where the *alignment θ*:

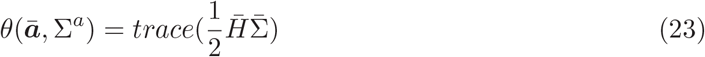

is a scalar that measures the relative orientation between (normalized) principal sensitivity vectors and (normalized) principal variability vectors. In other words, the more the directions of maximal variability 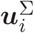 are aligned with the directions of maximal sensitivity 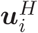 of the score, the larger the effect of variability on the *β* and hence on the expected action score.

In conclusion, the performance of a motor skill, or its expected action score can be approximated (and decomposed) as:

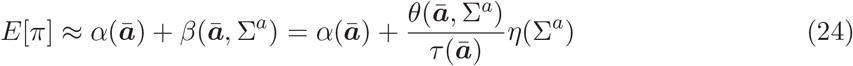

### 4.3 Application to throwing tasks

Throwing skills, as many other motor skills, are usually assessed by means of score functions which essentially define the objective of the throwing task. For instance, for a javelin thrower, the score may be a function of the longitudinal distance travelled by the javelin. The further the javelin lands, the larger the score assigned to the throwing action. Conversely, for a dart thrower, the goal is not to throw the dart as far as possible, but as accurate as possible, and hence, as in Fig 1A, the score function could assign a penalty increasing with the distance between the landing position of the projectile and the center of the target.

It should be noted that in our experimental protocol [21] participants did not receive any explicit performance feedback (or score) at end of each throwing trial (but they could see the arrival position of the ball on the target board) and therefore, in this work, in line with computational and experimental evidences [18, 28], we assume that participants optimize an accuracy score which penalizes the squared error between the outcome ***x*** of a release action and the target position ***x***_*T*_ [28]:

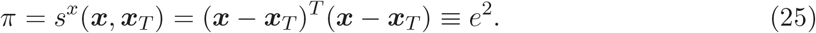

Written in this form, the accuracy score, represents a *task score s*^*x*^ : *R*^2^ → *R*^+^ which penalizes the two-dimensional action outcomes *x* with a scalar score *π*. To find the relationship between release actions and quadratic (task) error, i.e. the *action-to-score function s*^*a*^ : ***a*** ∈ *R*^6^ → *R*^+^, we need to express the task score (25) as a function of the release parameters.

Assuming a point-mass projectile and hence neglecting friction and Magnus’s forces, the projectile trajectory ***f*** (***a***, *t*) can be predicted from the release parameters ***a*** = ***p***_0_, ***v***_0_ as:

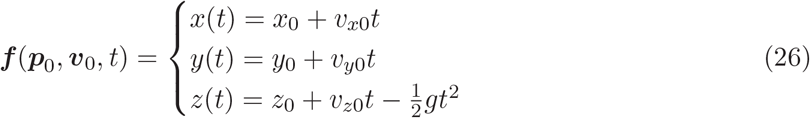

with *p*_0_ = [*x*_0_, *y*_0_, *z*_0_]^*T*^ and *v*_0_ = [*v*_*x*0_, *v*_*y*0_, *v*_*z*0_]^*T*^.

For a target board oriented as in Fig 1A, i.e. with the normal pointing in the longitudinal direction *y*, the time of impact of the ball with board can be estimated as 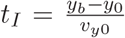, i.e. as the ratio between the longitudinal distance of the projectile with the board at release (*y*_*b*_ is the coordinate of the board with respect to the world frame) and the velocity of the ball along such direction (that is constant according to the model in (26)). Hence, at the time of impact, the projectile will hit the board at:

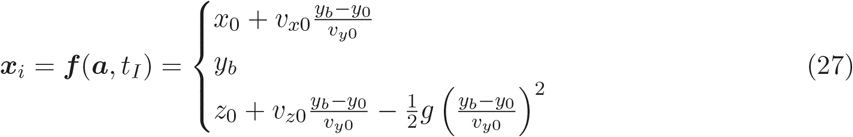

Substituting the above system of equations into (25), let us writing the *action score* as:

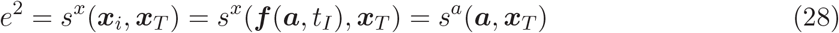

i.e, as a scalar function ***a*** ∈ *R*^6^ → *R*^+^ of the throwing action ***a***:

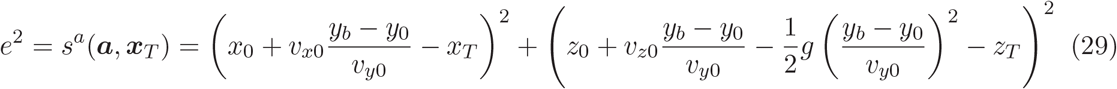

Given an individual release strategy with mean action ***ā***_0_ and covariance Σ^*a*^ and aiming at hitting a desired target ***x***_*T*_, the mean squared error (*E*[*e*^2^] = *E*[*π*]) can be approximated with (15):

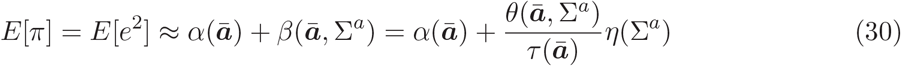

where *α*(***ā***) is just (29) evaluated at ***ā***, i.e. the quadratic error of the outcome of the mean action, and *β* can be decomposed into the three components *η, τ*, and *θ* by using the covariance matrix of the action strategy Σ^*a*^ and the 6 × 6 Hessian of (29) evaluated at ***ā***.

In this work, the 6×6 Hessian matrix of (29) is calculated with the Matlab Symbolic Toolbox for each target condition and for each individual strategy.

### 4.4 2D throwing simulation

To illustrate the Hessian-based decomposition in simple task with 2D actions, we considered a toy model of throwing. We assumed that a projectile is released from a fixed position and moves in a vertical plane to hit a target on a vertical axis at a given distance from the release position (Fig 3A). Thus, the action *a* corresponds to a 2D vector whose components are the horizontal release velocity 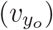 and the vertical release velocity 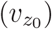, the outcome *x* corresponds to the arrival height on the vertical axis, and the score the squared distance from the target (*x*_*T*_).

We simulated distributions of release actions with a given mean ***ā*** and covariance Σ_*a*_ (Fig 3 and 4) by drawing 50 random samples from a multivariate Gaussian distribution. We simulated a target at a distance of 5 m from the release position and at height 0.7 m below it, reproducing the mean conditions for target T1 in our sample of 6D throwing participants (see below). In addition to decomposing the mean score according to equation (30) we performed TNC-Cost, UCM, and GEM analyses. For the TNC-Cost analysis, we followed the procedure described in [5]. For the UCM analysis and GEM analysis, we computed the Jacobian of the action-to-outcome function at the mean action and the orthogonal direction.

### 4.5 6D throwing experimental protocol and data analysis

Individual release strategies were obtained from the experimental dataset acquired in our previous study [21], where twenty right-handed participants (10 females, 10 males; age: 28.2 ± 6.8 years) performed a series of overarm throws, starting from a fixed initial position. All participants signed an informed consent form in accordance with the Declaration of Helsinki. The data collection was carried out in accordance with Italian laws and European Union regulations on experiments involving human participants. The protocol was approved by the Ethical Review Board of the Santa Lucia Foundation (Prot. CE/PROG.542). Participants were instructed to hit one of four circular targets arranged on a vertical target board placed at 6 m from the initial position (marked with a sign on the floor) and to start from a fixed posture (standing with the arm along the body). The four targets were custom made and consisted in white circles of 40 cm diameter, arranged on a rectangular layout on the target board. The distances between the centers were 70 cm vertically and 80 cm horizontally, similarly to Fig 1. Moreover, the targets midpoints in the horizontal direction were shifted with respect to the projected initial position of the participant: the left and right targets were centered respectively at 60 cm to the left and 20 cm to the right of the projected initial position of the throwers midline. An opto-electronic system (OptiTrack, NaturalPoint, Inc., Corvallis, OR, United States) operating at 120 Hz was used to capture whole-body kinematic information of the participants throwing actions and the corresponding ball trajectories.

For each trial and participant, the release action *a*, was obtained by fitting each of the three spatial components of the ball path with a 3rd-order polynomial function, and therefore the release position *p*_0_ and velocity *v*_0_ were obtained from the zero and first order coefficients, respectively. Then, this release action was used off-line in (27) and (25) to generate ‘ideal’ ball paths and scores, respectively, which were not influenced by friction and/or spinning effect of the ball. Trials in which the ball path did not intersect the target plane, or for which the ball was partially tracked by the optical system, were excluded from the analysis (13% of the total number of throws; we verified that differences in the number of trials across participants and conditions affect the values of some of the decomposition parameters only for sets with fewer trials than our smallest set). The error distribution, across trials, participants and target conditions, between experimental and ideal performance (mean squared error) is shown in Fig 9 (mean ± SD: 0.0012 ± 0.0912*m*^2^). The dataset is available in S2 Dataset.

**Figure 9.**
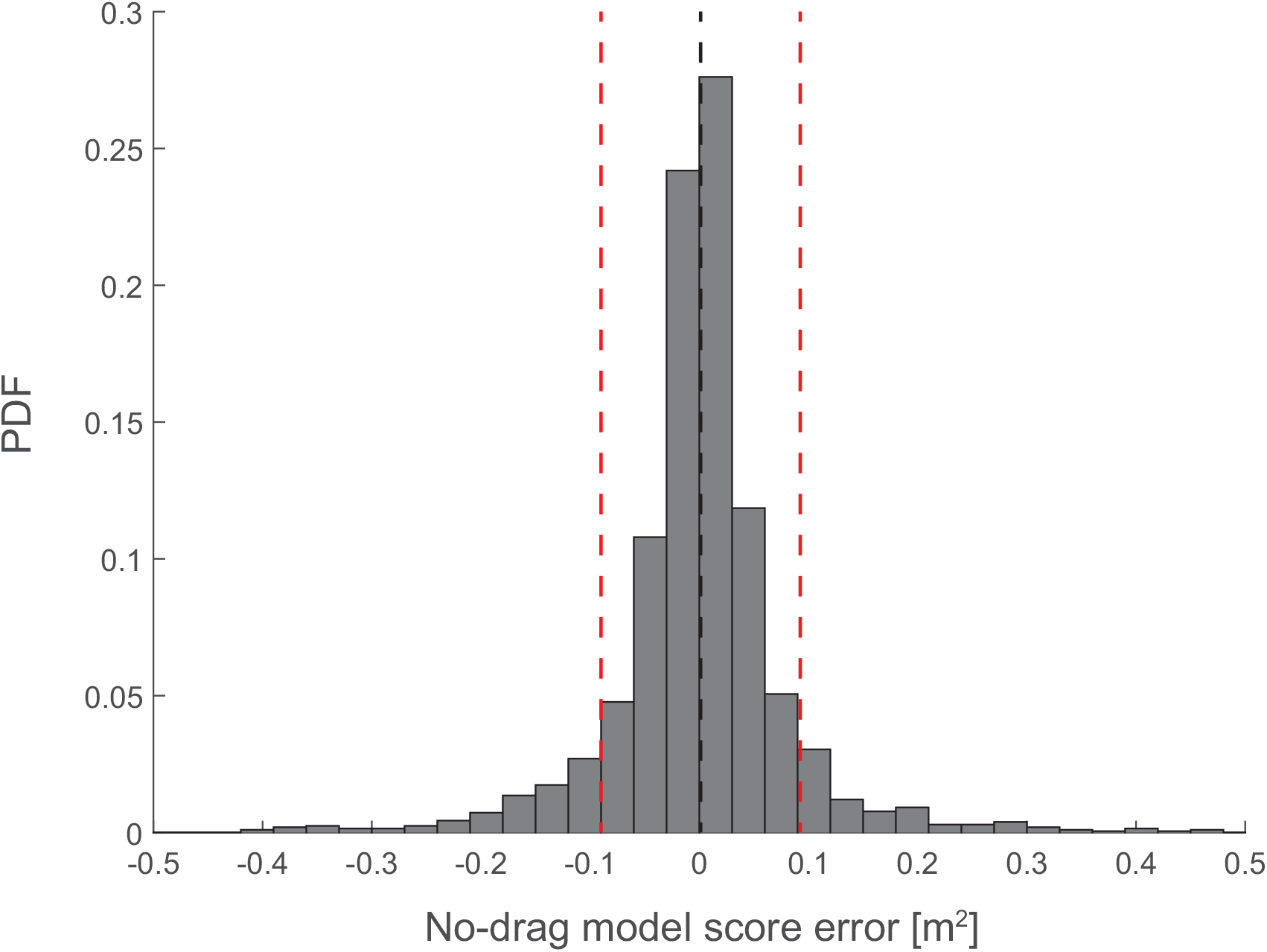
Error distribution between experimental score and score predicted with the no-drag model in equation (29). Black and red vertical dashed lines indicate mean and ±1*SD* of the distribution, respectively.

**Figure 10.**
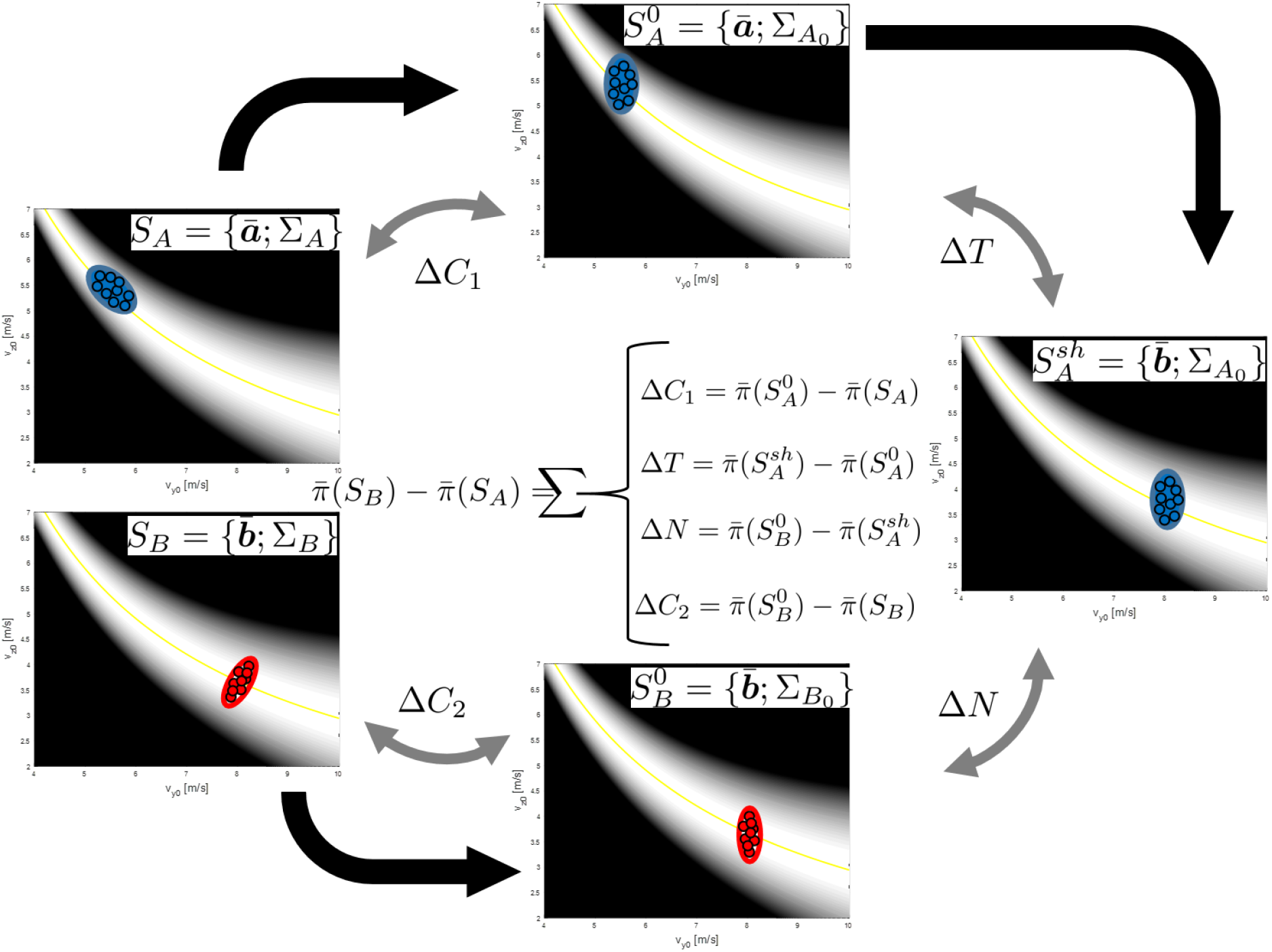
The TNC method proposed in [23]. Given two datasets, *A* and *B*, for instance the release strategies of two different partecipants, the method requires the generation of surrogate data-sets *S*^0^ (covariation-free) and *S*^*sh*^ (covariation-free but shifted mean). The difference in mean score between experimental and surrogate datasets are then used to calculate the relative tolerance Δ*T*, noise Δ*N* and covariation Δ*C*_1_, Δ*C*_2_ between the two strategies.

We next assessed the validity of the assumption that the individual action distribution is sufficiently localized around the mean action such that the second-order approximation of the sample mean score given in equation (29) as the sum *α* + *β* is adequate. To do so we computed the fraction of variance accounted for (VAF) by the approximation, defined as:

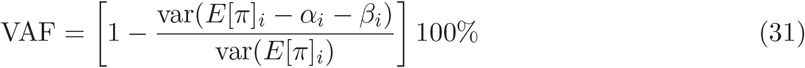

VAF is therefore defined as the variance of the error between the individual performance (sample mean score) and the second order approximation normalized by the variance of the population performance.

In summary, for each participant and for each target ***x***_*T*_, we estimated the mean-quadratic-error *E*[*π*], the mean action ***ā***, and the action covariance Σ^*a*^, with their respective sample mean and covariance: 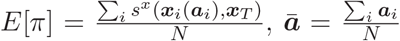, and 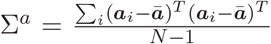, where *N* is the total number of successful actions, or trials, executed for target ***x***_*T*_.

## Statistical analysis

To assess the effect of target and participant identity on the score and on the indicators extracted from the Hessian-based decomposition, we fit a linear mixed-effects model (Matlab function *fitlme*) with target and participants identity (categorical variables) as fixed effects and participants intercept as random effect

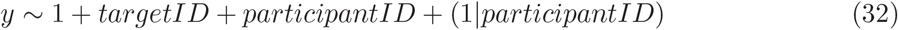

and we performed an analysis of variance for linear mixed-effects model (Matlab function *anova*), thus taking into account the repeated measures design.

## 5 Appendix

### A. Relation to TNC approaches

#### A.1 Relation to Müller & Sternad 2004

In this section we show that, when the score function is smooth, and the action distribution is sufficiently localized, it is possible to derive (Hessian-based) analytic expressions to isolate the three components of the TNC approach by Müller and Sternad [23], shown in Fig.10. Given two experimental strategies, such as *S*_*A*_ and *S*_*B*_ in the figure, the TNC method requires the generation of surrogate data-sets, 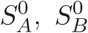 and 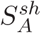, to decompose the difference in expected score 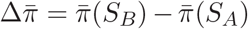, into the sum of four independent components: Δ*C*_1_, or *covariation*, is the difference in expected score between the strategy *S*_*A*_ and the (surrogate) strategy 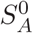, that is obtained by removing (via random permutations) any linear/non-linear correlation between the variables of the data-set *S*_*A*_. Similarly, for *S*_*B*_, a surrogate uncorrelated data-set 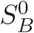 is used to quantify the difference in performance Δ*C*_2_ due to covariations in *S*_*B*_. Notice that 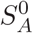 and 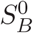 have the same mean (***ā*** and 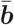, respectively) as their original data-sets and only differ with respect to their original data-sets in terms of variability. A third surrogate data-set 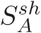 is generated by *shifting* the location of 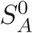 (i.e. ***ā***) to the average location 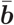 of the *S*_*B*_ data-set. The *tolerance* component is hence quantified as 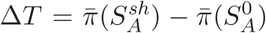, has the two data-sets have same variability and differ only in terms of their average location. Lastly, the *noise* component is extracted as the difference in average performance between the surrogate data-set 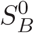 and 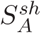, i.e. 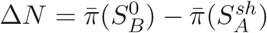.

Assuming that motor strategies are drawn from a localized distributions, *S*_*A*_ = {***ā***; Σ_*A*_} and 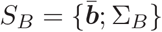, our method allows the estimation of all four components without using surrogate data-sets and random permutation. In this case, the covariance matrices 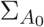 and 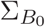 of the uncorrelated strategies 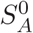 and 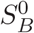, can be simply computed as diag(Σ_*A*_) and diag(Σ_*B*_), i.e. as the matrix of the diagonal elements (variances) of Σ_*A*_ and Σ_*B*_, respectively. Knowing the Hessian matrix *H*_***ā***_ and 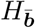 at the two locations ***ā*** and 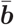, respectively, allows to approximate Δ*C*_1_,Δ*T*, Δ*N* and Δ*C*_2_, simply as:

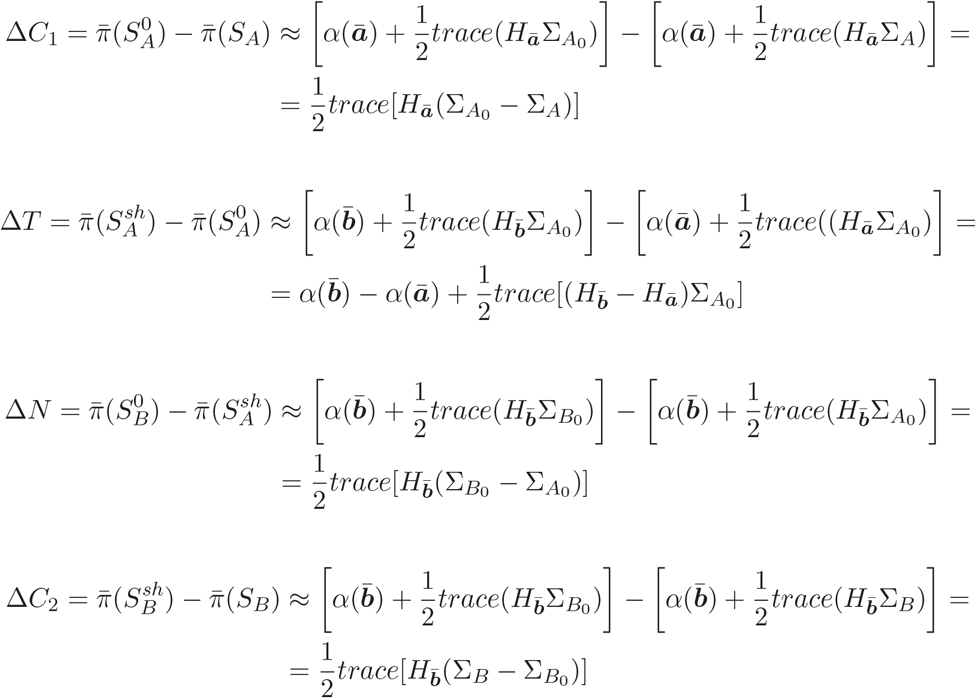

Hence, it follows that the difference in expected performance between the two strategies can also be approximated as:

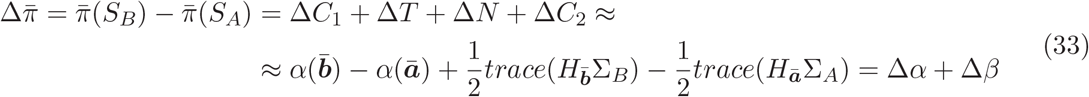

#### A.2 Relation to Cohen & Sternad 2009

An alternative TNC approach, the TNC-Cost, has been proposed by Cohen & Sternad [5] to overcome some of the limitations of the original TNC described in the Appendix A1. In the novel approach, the parameters of the decomposition are no longer dependent on the sequence of calculation, and the components are quantified with respect to an optimized data sets derived by transformation of one of the features.

*The T-Cost*. The T-cost is the algebraic difference between the mean score of the experimental dataset and the mean score of an optimized dataset that differs from the first dataset only in terms of the mean action. More specifically, the action space is firstly discretized in a grid of *n* points that are used to shift the action distribution of the original dataset. Hence, for each grid point the original and the shifted dataset have the same action variability and only differs with respect to the mean action. For each grid points the method evaluates the mean score of each shifted dataset, such that the optimal one is the one which results in the minimal (i.e. best) mean score.

In the assumption that experimental dataset *S*_*A*_ has a small dispersion, we can apply our framework to calculate, analytically, the T-cost. In particular, let ***ā*** and Σ_*A*_ be the mean action and action covariance of the experimental dataset *S*_*A*_, and let ***ā***^*^ be the shifted mean of the optimized data set 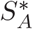, then the T-Cost can approximated as:

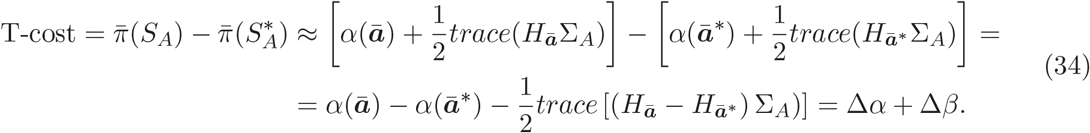

In the hypothesis that the original dataset had already an optimal mean action, i.e. on the solution manifold, and that the grid was sufficiently discretized such that also the optimal action belongs to the solution manifold, than the tolerance cost of Cohen & Sternad would correspond to a Δ*β* due to differences in the local geometry of the score between the experimental and the optimized datasets, i.e *H*(***ā***) ≠ *H*(***ā***^*^).

Notice however, that the tolerance cost depends on the action covariance of the original dataset; hence it does not provide a unique measure of *tolerance*/*sensitivity* to errors of the score function. In other words, given two experimental datasets *S*_*A*_ and *S*_*B*_ with the same mean action 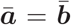 but different covariances, one would measure a different Tolerance-cost for the two distributions, despite locally, the score has the same Hessians for both distributions, i.e. 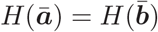.

*The N-Cost*. The N-cost is the algebraic difference between the mean score of the experimental dataset and the mean score of a dataset with an optimized *noise*. More specifically, the optimal noise is obtained through a sequence of *n* steps which progressively shrunk the data points (the actions) of the original dataset towards its mean action. For each step, the mean score of the shrunk dataset is calculated and the optimal dataset is identified as the one with minimal mean score.

In the assumption that experimental dataset *S*_*A*_ has a small dispersion, we can apply our framework to calculate, analytically, the N-cost:

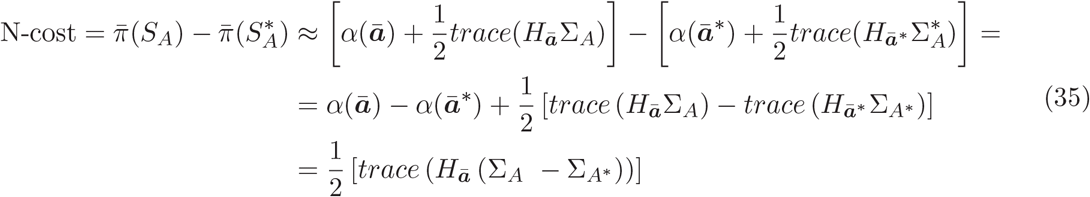

where in the last identity we have used the fact that the experimental and optimized data-set have the same mean action, hence the same *α* and Hessian. This result shows that in general the N-cost depends on both the local tolerance of the score and the difference in covariance between the experimental and the optimized data-set. In the particular case in which the shrinking procedure returns an optimal dataset which is just a single point [5], i.e. Σ_*A*\*_ = 0, the N-cost corresponds to our *β*, highlighting the fact that this parameter does not in general provides a clear description of *noise* (meant as action variability) given that it mixes all the contributions that are due to the action variability and the geometry of the score.

*The C-Cost*. The C-cost is the algebraic difference between the mean score of the experimental dataset and the mean score of an optimized dataset that differs from the first only in terms of correlation between action variables, hence having the same mean action and uncorrelated variability of the original dataset. Following our framework, in the hypothesis of localized distribution and smooth score function, the C-cost, or the difference between the original and the optimized dataset becomes:

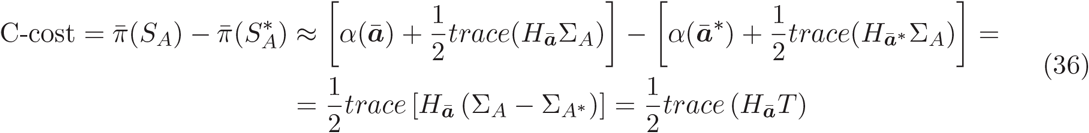

where the *T* matrix has all zeros on the main diagonal given that the two datasets have the same uncorrelated noise.

### B. Examples of individual distributions of release parameters in unconstrained 6D throwing

When actions are high-dimensional, as in unconstrained throwing where the action is a six-dimensional (6D) vector, it is impossible to visualize both the score and the individual strategies in a single plot, as in the 2D throwing case. Fig 11 shows the action score and individual throwing strategies in terms of pairs of release parameters (9 of the 15 possible pairs of 3 position and 3 velocity variables; *rows*) for five exemplary participants (*columns*) throwing at target T1. The last three *rows* are also illustrated in Fig 5 in the main text. Participants have been sorted from left to right according to their average release speed, hence *P*10 is the slowest thrower and *P*11 the fastest and they have similar horizontal and vertical release velocities as the five simulated 2D strategies illustrated in Fig 3B-C in the main text. The gray-scale shaded contours indicate the score associated to each pair of release parameters, i.e. the squared distance between the ball arrival position on the vertical target plane and the center of the target. The domain of each position and velocity variable corresponds to the population mean ± 3 standard deviations, while all the remaining release parameters are considered constant and fixed to the subject-specific mean action. There are 15 different gray-scale levels: the white area defines actions which have a score smaller or equal to 0.04 *m*^2^, i.e. actions that land inside the target, which has radius 0.2 *m*. Notice that the regions corresponding to actions with the same score (i.e. same gray shading) have different shapes (or geometry) across planes and participants, as they depend on the individual mean release action. The wider the white areas around the mean release parameter, the more tolerant is the action-to-score function to stochastic perturbations. For instance, in the 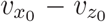 plane (*8-th row*), the lowest penalty (white) area looks like an ellipse, whose orientation suggests that the 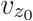 direction is less tolerant/more sensitive to action variability and whose size, increasing with release speed, suggests that tolerance is higher for higher speed. In general, position variables appear more tolerant than velocity variables and the shapes of the white regions in the 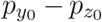 plane (*third row*), in the same-axis position-velocity planes (*fourth to sixth rows*), and in the 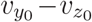 plane (*ninth row*) indicate that release variables corresponding to the lowest score (throws hitting the target) are negatively correlated.

**Figure 11.**
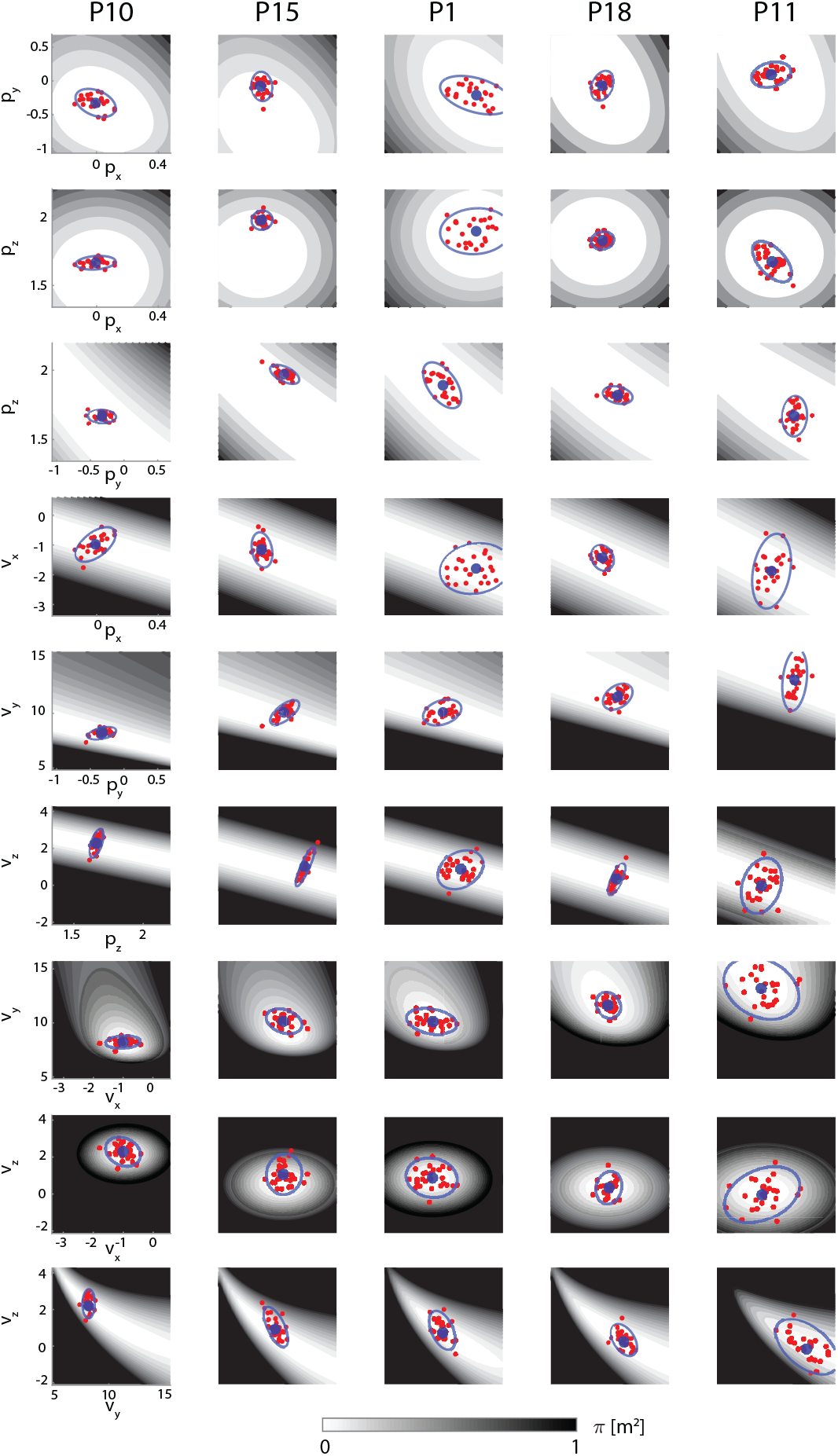
Examples of distribution of release position and velocity for five representative participants (target T1). Each row illustrates a pairs of release parameters, including all combinations of position variables (*p*_*x*_-*p*_*y*_, *p*_*x*_-*p*_*z*_, *p*_*y*_-*p*_*z*_), all combinations of position and velocity variables for each axis (*p*_*x*_-*v*_*x*_, *p*_*y*_-*v*_*y*_, *p*_*z*_-*v*_*z*_), and all combinations of velocity variables (*v*_*x*_-*v*_*y*_, *v*_*x*_-*v*_*z*_, *v*_*y*_-*v*_*z*_). Red circles represent release parameters of individual throws. Blue circles and ellipses represent mean and covariance (two standard deviations) of each parameter distribution. The gray level map shows the local score, as a function of the release parameters, underlying each throwing strategy. Note that differences in the background geometry of the maps reflects individual differences in the average action (mean position and velocity vectors at release). The wider the white area around the mean action, the more tolerant the score is to stochastic perturbations. Participants have been sorted from left to right according to their average release speed.

The distributions of the release parameters (*red circles*), summarized in each plot of Fig 11 in terms of mean (*blue circle*) and covariance (two-standard deviations, *blue ellipse*), also differs remarkably across planes and participants. For instance, *P*11 (*fifth column*), on average, releases the ball with a slower vertical velocity 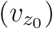, compared to other participants, such as *P*10 (*first column*) and *P*1 (*third column*), who instead throw with higher longitudinal velocity 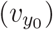. The amount of variability in the 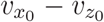 (*eight row*) and 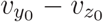(*ninth row*) planes also differs between the same three participants, as *P*11 shows a much wider covariance ellipses than *P*1 who, in turn, shows a wider covariance ellipses than *P*10. However, interestingly, their performance is similar, as the large variability of *P*11 in the 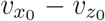 and 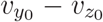 planes is partially compensated by being in a more tolerant region of the action score.

Also to notice are different patterns of covariance/correlation across individual strategies. For instance, *P*10, who is the least variable participant, shows no correlation between 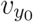 and 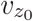, while *P*11, who is the most variable participant, shows a negative correlation: reducing the vertical release velocity proportionally to an increase in the longitudinal release velocity. This allows *P*11 to remain in the lowest penalty region as much as *P*1 and *P*10 and hence to have a similar mean score. Notice that *P*15 has a mean release velocity located close to the edge of the white region and hence on average does not hit the center of the target. However, *P*15 has a mean score similar to those of *P*10, *P*1, and *P*11, in part due to the small and, in most cases, well aligned covariance ellipses. Finally, *P*18, the best performing participant, shows mean release parameters at the center of all white regions and small and well aligned covariance ellipses.

In sum, the examination of the distribution of several pairs of release action parameters and the associated score suggests that individual throwing strategies differs in terms of mean action, action variability, and relationship between action variability and geometry of the action-to-score function. However, as the distribution is 6-dimensional and there are 15 different pairs of action variables, it is not possible to identify by visual inspection a unique source of the inter-individual differences in throwing strategies and to systematically explain the relationship between action distribution and throwing performance. These limitations can be overcome by introducing the Hessian-based decomposition of the mean score that we developed to provide a compact and informative description of the key features of the action distribution characterizing individual strategies and directly related to performance.

### C. Score sensitivity (Hessian) and action variability (covariance) in unconstrained 6D throwing

The Hessian-based decomposition allows to characterize the structure of individual variability and its relation with the local geometry of the score, overcoming the limitations of a qualitative description of the action distribution and the score function, which is challenging when the score is defined over a high-dimensional space of action variables. Similar to the UCM and GEM method, which use Jacobian matrices to split motor variability along task-relevant and task-irrelevant directions, we use the Hessian matrix to quantify *score-relevant* variability affecting the mean score. Here, we illustrate how, across participants, a few eigenvectors of the Hessian matrix (principal sensitivity directions) associated with the largest eigenvalues identify the score-relevant directions that determine whether action variability affects the mean score or not. We characterize the structure of the Hessian and the action covariance matrices, which define the terms of the Hessian-based decomposition.

Fig 12A shows the distributions, across participants and for each target, of the eigenvalues or singular values of the Hessian matrix. Because in our throwing task the outcome space is two-dimensional, the solution manifold is a four-dimensional manifold embedded in the six-dimensional action space (see Appendix C). Hence, on the solution manifold, the Hessian matrix will only have two non-zero singular values, whose associated eigenvectors or singular vectors defines a score-relevant plane. The corresponding eigenvalues quantify the sensitivity of the score function along that direction. Away from the solution manifold however, the Hessian matrix is also influenced by the non-linearities introduced by the mapping between actions and outcomes. Hence, for participants that do not have optimal mean actions, the Hessian matrix can have additional singular values which are different from zero. Fig 12A shows, however, that the contribution of the third and fourth singular value is negligible compared to the first two. Finally, focusing on the first two singular values, we notice that the sensitivity is slightly anisotropic, with the first sensitivity, on average, about 10% higher than the second. Furthermore, the first sensitivity shows the largest variability across participants and target conditions.

**Figure 12.**
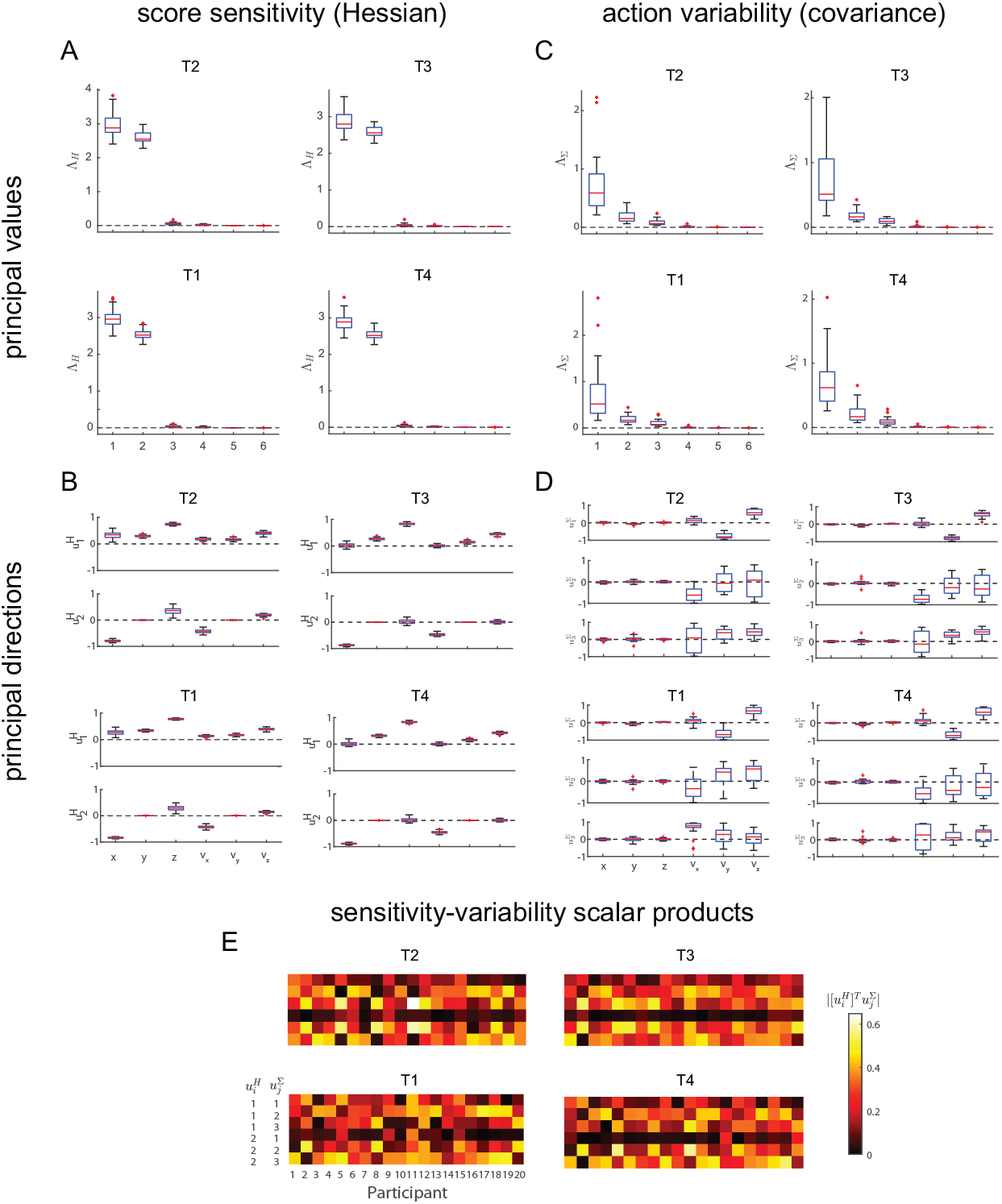
Sensitivity of the action-to-score function (Hessian matrix), action variability (covariance matrix), and their relationship across participants. (A) Distributions of the Hessian eigenvalues across participants for each target. In our scenario, the outcome space is two-dimensional and the solution manifold is a four-dimensional manifold embedded in the six-dimensional action space (see Appendix S1 Appendix C). Hence, the local tolerance of each participant is dominated by the first two eigenvalues of the Hessian matrix. (B) Distributions of the first 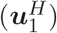 and second 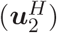 principal sensitivity directions across participants for each target. The first principal sensitivity direction is dominated by the vertical release position and velocities while, the lateral and longitudinal components contributes ‘equally’ for target T1 and target T2, while for target T3 and target T4, the longitudinal components were ‘more score relevant’ than the lateral ones. The second principal sensitivity direction is instead dominated by the lateral release position and velocity. (C) Distributions of the eigenvalues of the covariance matrix across participants for each target. The first three principal components explain 95% of the total variation. (D) Distributions of the first 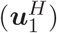, second 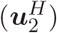, and third 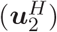 principal variability directions across participants for each target. (E) Absolute values of the scalar products between all pairs (*i, j*) of the first two principal sensitivity directions 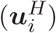 and the first three principal variability directions 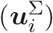.

The two principal sensitivity directions 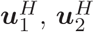 define locally, i.e. around the mean action, a *sensitivity plane* embedded in the six-dimensional action space of the release parameters. Fig 12B shows the distributions of the principal sensitivity directions across participants and for each target. Across targets, the first principal sensitivity direction 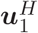 is dominated by the vertical components (both position and velocity) of the release parameters and the second principal sensitivity direction is instead dominated by the lateral components (*x*_0_, *v*_*x*0_) of the release parameters.

In terms of action variability, for all participants and for all target conditions three principal components were able to explain 95% of the total variation, as shown in Fig 12C. Across participants and target conditions, the eigenvalue of the first principal component shows large variability across participants. It should be noted that the covariance matrix and, thus, the number of principal components is coordinate-dependent and in our scenario the action vectors contain both position and velocity variables which have different units. To assess the robustness of the estimation of the dimensionality of the action variability, we performed a principal component analysis on the correlation matrix rather than on the covariance matrix. The analysis of the eigenvalues of the correlation matrix confirmed that across participants and target conditions there were no more than three eigenvalues grater than one [17], supporting the conclusion that three components are sufficient to adequately describe the action variability.

Fig 12D illustrates the distribution of the first three principal variability directions 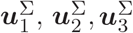, i.e. the eigenvectors of the covariance matrix associated to the three largest eigenvalues, indicating the directions along which most of the variation occurs. As for the principal sensitivity directions, a number of features of the principal variability directions are consistent across targets. For all four targets, the first direction captures a negative correlation between longitudinal (*v*_*y*_) and vertical (*v*_*z*_) release velocities and the second direction is dominated by the lateral velocity (*v*_*x*_) but with variable contribution of the other velocity components. The distribution of all three directions, in larger measure for the second and third direction, however, are broader than those for the principal sensitivity directions, indicating that there are larger inter-individual differences in the structure of the action variability than in the sensitivity of the score around the mean action.

The distributions of principal sensitivity directions and principal variability directions illustrated in Fig 12B and D characterize features of the throwing strategy consistent across targets and participants. However, it is the selection of specific directions and, even more, their geometric relationship that determines the performance of individual participants. For example, two strategies with identical mean action, which implies identical sensitivity, may have different mean scores because of different alignments of the principal variability directions with respect to the principal sensitivity directions. Fig 12E shows the absolute values of the scalar products between the 6 pairs of principal sensitivity directions 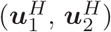 and principal variability directions 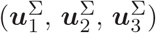 in all 20 participants. Such complex pattern of scalar products between principal sensitivity and variability directions highlights a remarkable inter-individual variability in the relationship between action variability and score sensitivity.

### D. Coordinate invariance

Approaches based on covariance matrices for the analysis of variability, such as the UCM, have often been criticized for their dependence on the choice of coordinates. Similar critiques have also been highlighted for the TNC approach that does not use (directly) covariance matrices: ”for instance, one can always rotate the frame of reference to get variables that have zero covariance” [29]. Furthermore, it is well known that Principal Component Analysis is sensitive to co-ordinates, especially when the multivariate data contains variables with different units. For instance, in this work the action vector contains position that are measured in meters and velocities that are measured in *ms*^−1^. Should we rescale the action space to have comparable variances between positions and velocities? Would scaling affect our results? Here we show that this is not the case and that both ours and the TNC approach [23] are invariant under affine coordinate transformations. In fact, scaling, rotations and translations, i.e. any affine transformation of the action space, does not only affect covariance matrices (and hence correlations among variables) but also affects the performance manifold, in particular the structure of its Hessian.

Let’s assume that we have two sets of coordinates {*a*} and {*b*} with which we can parameterize the *n*-dimensional action space 𝒜 ⊂ℝ^*n*^ and that the map *g* describe the relationship between the two coordinate system:

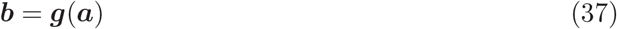

when the map is non-linear, its first-order approximation, around a point ***ā***.can be expressed as:

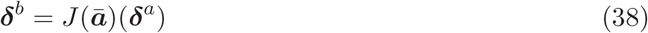

where 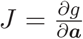 is the *n* × *n* Jacobian matrix evaluated at ***ā***.

The score *π* is a scalar and therefore does not dependent on the choice of coordinates used to express the score function. In both coordinate systems we can write:

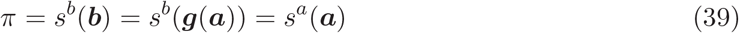

By differentiating the last equality with respect to the {*a*} co-ordinates, we find a well-known expression between the gradients in the two different co-ordinate systems:

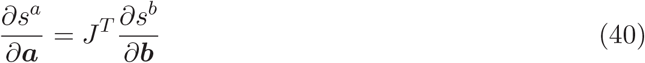

Differentiating again the above expression we can express the Hessian of the score in the two coordinate systems:

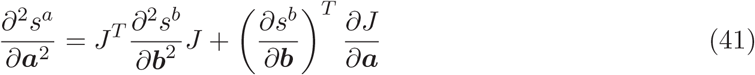

Hence:

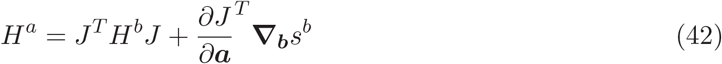

When the average action belongs to the solution manifold (**∇**_*b*_*s*^*b*^ = 0), or when the change of co-ordinate is affine 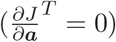, the second term on the right hand-side is zero. In this case, the Hessian is a tensor and *β* does not depend on the co-ordinates used to parameterise the action space. In fact, let Σ^*b*^, be the covariance of the action expressed in the {*b*} coordinates, then, if the distribution is localized (small variability), the covariance in the {*a*} co-ordinates can be estimated as:

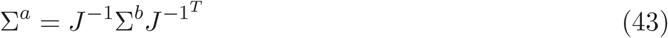

and hence:

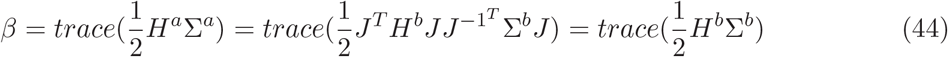

where we have used the cyclic properties of the trace (*trace*(*ABC*) = *trace*(*CAB*)) to simplify the last equality.

Conversely, for highly non-linear change of coordinates, or for average actions that are ‘far’ from the solution manifold, the second term on the right-hand side of (42) may not be negligible. In such case, the Hessian looses its tensorial property, and *β* becomes a co-ordinate dependent measure:

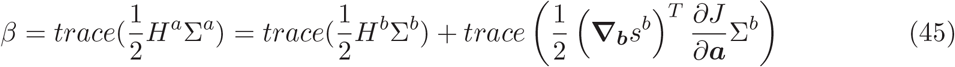

### E. Non-zero Hessian’s eigenvalues in the presence of redundant actions

In human motor control, the map between action and task variables represents a ‘change of co-ordinates’ ***x*** = ***f*** (***a***), which often is non-linear and redundant. This latter properties of the map, makes the Jacobian 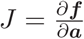 a rectangular matrix with *n* columns (dimension of the action space) and *m* rows (dimension of the task space). In this case, equation (42) becomes:

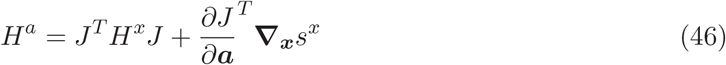

where *H*^*x*^ and **∇**_*x*_*s*^*x*^ are the *m* × *m* Hessian and the *m* × 1 gradient of the task-score function, respectively, and *H*^*a*^ is the *n* × *n* Hessian of the action-score function. Again, either on the solution manifold, or for a linear map between actions and outcomes, the second term on the right-hand side disappears, and *H*^*a*^ will only have *m* < *n* non-zero eigenvalues. Conversely, away from the solution manifold and for highly non-linear change of co-ordinates, the term 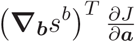 will in general affect the number of non-zero eigenvalues, as well as the symmetry and positive-definiteness of the Hessian matrix.

## Acknowledgments

We thank Etienne Burdet for reading the manuscript and providing insightful comments and many suggestions and Maura Mezzetti for helping with the statistical analysis.

